# Na_V_1.7 targeted fluorescence imaging agents for nerve identification during intraoperative procedures

**DOI:** 10.1101/2024.04.06.588368

**Authors:** Junior Gonzales, Dauren Adilbay, Paula Demetrio de Souza Franca, Raik Artschwager, Chun Yuen Chow, Tara Viray, Delissa S. Johnson, Yan Jiang, Snehal G. Patel, Ian Ganly, Christina I. Schroeder, Jason S. Lewis, Glenn F. King, Thomas Reiner, Nagavarakishore Pillarsetty

## Abstract

Surgeries and trauma result in traumatic and iatrogenic nerve damage that can result in a debilitating condition that approximately affects 189 million individuals worldwide. The risk of nerve injury during oncologic surgery is increased due to tumors displacing normal nerve location, blood turbidity, and past surgical procedures, which complicate even an experienced surgeon’s ability to precisely locate vital nerves. Unfortunately, there is a glaring absence of contrast agents to assist surgeons in safeguarding vital nerves. To address this unmet clinical need, we leveraged the abundant expression of the voltage-gated sodium channel 1.7 (Na_V_1.7) as an intraoperative marker to access peripheral nerves *in vivo*, and visualized nerves for surgical guidance using a fluorescently-tagged version of a potent Na_V_1.7-targeted peptide, Tsp1a, derived from a Peruvian tarantula. We characterized the expression of Na_V_1.7 in sensory and motor peripheral nerves across mouse, primate, and human specimens and demonstrated universal expression. We synthesized and characterized a total of 10 fluorescently labeled Tsp1a-peptide conjugates to delineate nerves. We tested the ability of these peptide-conjugates to specifically accumulate in mouse nerves with a high signal-to-noise ratio *in vivo*. Using the best-performing candidate, Tsp1a-IR800, we performed thyroidectomies in non-human primates and demonstrated successful demarcation of the recurrent laryngeal and vagus nerves, which are commonly subjected to irreversible damage. The ability of Tsp1a to enhance nerve contrast during surgery provides opportunities to minimize nerve damage and revolutionize standards of care across various surgical specialties.

## INTRODUCTION

Nerves are fibrous structures present throughout the human body and they are essential for sensory and motor function. Every surgical procedure, regardless of location and complexity, possesses nerve damage risk. Since there are no FDA-approved agents to visualize nerves, identification must rely on anatomical landmarks and surgeon experience. Surgical patients can experience iatrogenic nerve damage (IND) caused intentionally or inadvertently from surgical manipulations that include cutting, ligating, thermal cauterization, or blunt trauma.^1–6^ About 25% of the 610,000 irreversible nerve injuries that occurred in 2024 were directly attributed to surgical interventions.^6,78–11^ Oncologic surgeries pose significantly increased risk of peripheral nerve damage because of changes in tissue texture, altered anatomy due to malignant lesions, and the necessity for complete tumor resection, which can lead to disability in patients.^12,13^ For example, in head and neck surgery, damage to the cranial nerves can result in facial disfigurement, voice hoarseness, respiratory distress, and dysphagia.^14,15^ Prostatectomies are often associated with a notable risk of nerve damage, resulting in erectile dysfunction, infertility, and penile nerve amputations.^16,17^ Similarly, other oncologic surgeries, including those for breast cancer, abdominal malignancies, and extremity soft tissue tumors, also put nerves at risk of injury.^18–20^ The limitations of current surgical instruments in the operating room (OR) pose significant concerns regarding nerve injury prevention, especially in robot-assisted and laparoscopic surgeries where exposing and localizing nerve structures remains challenging.^1,2,21^ Notably, the recurrent laryngeal, facial, vagus, brachial plexus, ulnar, sciatic, peroneal and other similar nerves are highly susceptible to damage during various common surgical interventions, including thyroid surgery, parotidectomies, carpal tunnel syndrome treatment, varicose vein procedures, arthrodesis, and lymph node neck dissections.^22,23^

In the OR, reducing iatrogenic complications requires an operating surgeon (OS) to possess awareness of the precise locations of nerves. For instance, nerves can be mistakenly identified as vessels, tendons, or other scarred tissues that are deeply embedded in the surrounding tissues.^24–27^ Current approaches for pre-operative assessment of nerve location include magnetic resonance imaging (MRI), computed tomography (CT), and ultrasound guidance; however, these modalities are not applicable for identifying all nerves. This prompts physicians to primarily rely on their anatomical knowledge, subjective assessments, intuition, and experience to identify and preserve nerves during critical surgeries, thereby increasing the risk of IND.^28–30^

While intraoperative technologies exist to improve planning and preservation for larger nerve bundles,^31–34^ a detailed delineation or localization of peripheral nerves remains challenging.^13^ In the OR, surgeons primarily rely on intraoperative tools such as magnification loupes, stationary microscopes, and electromyography electrodes for nerve identification.^35,36^ However, these tools require the identification or exposure of nerves, or proximity to the tissue surface for stimulation, rendering them unsuitable for all types of interventions.^37^ The lack of accurate techniques for nerve identification may lead to IND, resulting in long-term impairments for the patients. Currently, nerve fluorescence imaging (NFI) is on a trajectory to become a GPS of the operating theater. Gibbs and his colleagues constructed a library of neuro-specific fluorescent agents using red oxazine fluorophores to visualize peripheral nerves.^38^ Subsequently, Frangioni et al. introduced systemic nerve-targeted contrast agents, BMB and GE3082, which light up myelinated nerves after a single injection.^39^ In addition, Nguyen’s group developed a nerve peptide (NP41) that binds to nerve-associated connective tissues.^40^

NFI surgery along with the creation of a fluorescent nerve-targeting agent would provide an opportunity to significantly reduce IND. Despite the clear medical need and its importance for those undergoing surgery as a standard of care, peripheral nerves have proven to be a difficult target to study, not only attributed to the molecular inaccessibility, natural unique/non-specific structure, and the exquisite potentially targeting of their functional proteins, but once perturbed/damaged, nerves degrade rapidly.^38,41–43^

To address this challenge, we turned our attention to the voltage-gated sodium channel, subtype 1.7 (Na_V_1.7) that is expressed on peripheral nerves. Natural compounds have proven to be excellent pharmacological tools for addressing medical challenges, including targeting hard-to-drug targets such as Na_V_1.7. ^44–49^ We took advantage of the highly specific, serum-stable disulfide-rich Nav1.7-targeting peptide Tsp1a, isolated from the venom of a Peruvian tarantula, to develop nerve imaging agents.^43,50,51^ First, we conducted extensive *ex vivo* characterization of human, mouse and primate nerves obtained from cadavers and/or freshly excised samples to demonstrate universal Na_V_1.7 expression. We prepared a library of ten Tsp1a tracers for delivering red and near-infrared (NIR) fluorescence sensors to peripheral nerves for *in vivo* imaging. The clinical agents, all of which included a Tsp1a peptide core, were designed based on the following considerations: reproducible chemical manufacturing, durability, the optical range of clinically available surgical cameras, the effect of chemical modification on Nav1.7 affinity, deep tissue penetrability, minimal photobleaching and potential surgical simulation.^52–54^ We manufactured two Tsp1a scaffolds, Tsp1a-K4 and Tsp1a-Pra0, in which either a lysine residue (K4) or a propargyl group (Pra0) were used as attachment sites for NIR-sensors. All of the conjugates but one retained nanomolar potency for inhibition of human Na_V_1.7. Tsp1a-Pra0 conjugated to the FDA-approved IR800 fluorophore exhibited high chemical purity, excellent yield, high affinity for Na_V_1.7, stellar penetration characteristics, and stable fluorescence over time. Therefore, it was chosen as the lead candidate for pharmacokinetic studies and visualization of nerves during thyroidectomy in non-human primates (NHPs).^55^

During thyroidectomy, the left recurrent laryngeal nerve and vagus nerve were highlighted with excellent fluorescence signal *in vivo* using our tracer in NHPs. By repurposing Tsp1a peptides for fluorescence-guided surgery, we showed that the Tsp1a-IR800_P_ tracer is clinically relevant and able to help when delineating nerves during intraoperative interventions. Since the technology/tool is immediately available, quantifiable and has the potential to directly impact the surgical standard of care by lending contrast to nerves, it already surpasses any current technique for nerve identification in a clinical setting.

## RESULTS

### Neuronal expression of Na_V_1.7 is conserved across species

To delineate the extent and distribution of Na_V_1.7 expression in humans (Figure 1a), we dissected clinically relevant nerves from six diseased cadavers (n = 3 males and n = 3 females). When analyzing the expression of Na_V_1.7 receptors with anti-Na_V_1.7 antibody immunohistochemistry (IHC) in nerves from the extremities, differences in Na_V_1.7 expression were found between genders (average expression in males = 21.4 ± 13.1%, and females = 20.5 ± 8.9%, p = 0.0378, Figure 1b). To understand the natural anatomical distribution of Na_V_1.7 in humans, we reported the Na_V_1.7 expression as it is and we created a rubric of Na_V_1.7 expression (Figure 1c-f, S1a, n = 5 repeated adjacent nerve). The nerves and organs were subsequently divided into no, low, medium, high, and very high expression, according to the abundance of Na_V_1.7 in the tissue. Based on the level of expression of Na_V_1.7, the tissues and nerves were categorized into five groups. All analyzed nerves had Na_V_1.7 expression ranging from medium (10–15%) to high (15– 20%) or very high (>20%) levels. Other surrounding tissues, such as arteries, veins, muscles, and tendons had low or no expression of NaV1.7 (<5%); the difference was statistically significant.

**Figure 1.**
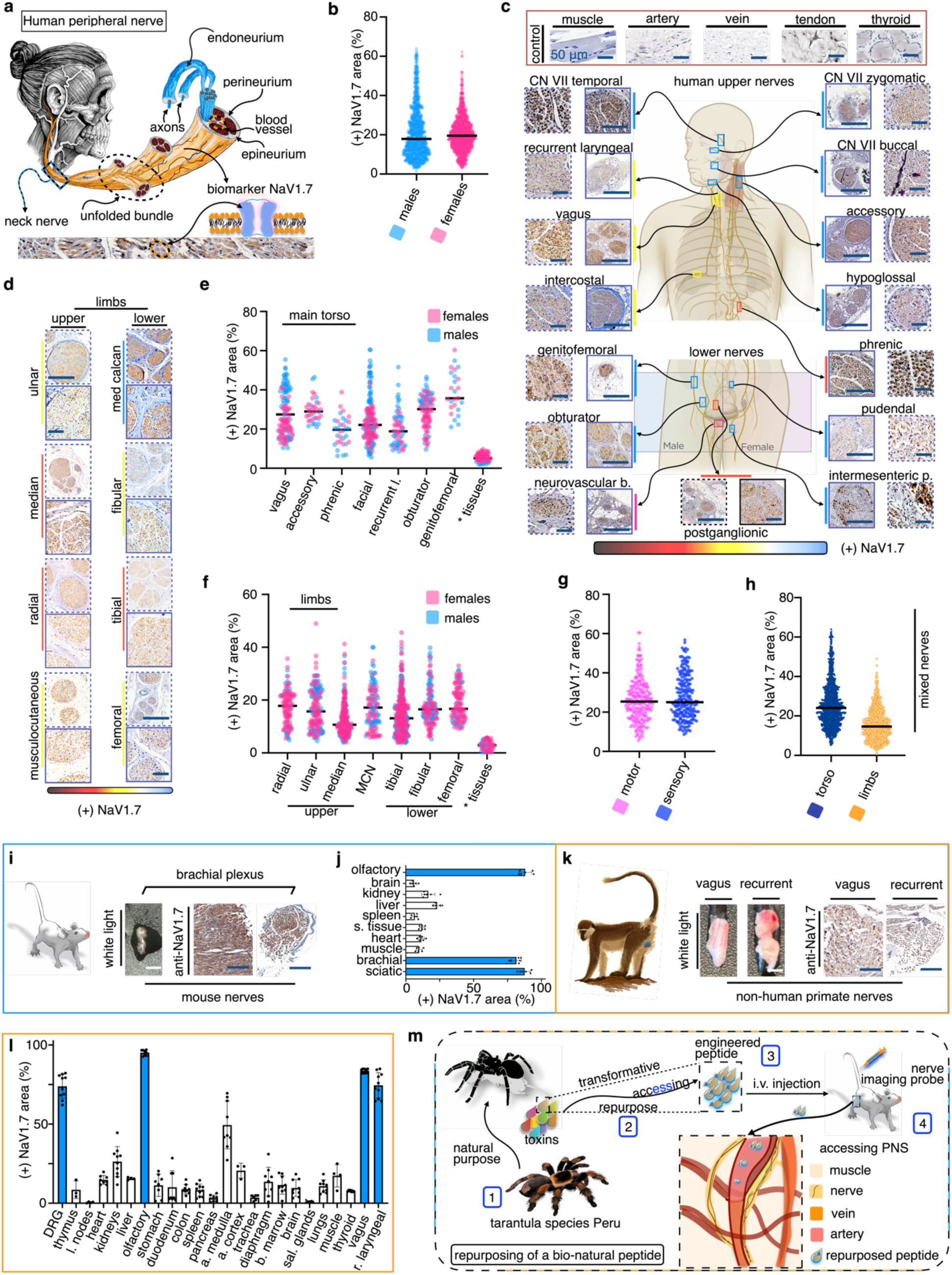
Expression of Na_V_1.7 in the peripheral nerves of humans, mice and NHPs. (a) Schematic illustrating the general architecture of a peripheral nerve. Individual nerve fibers (axons) are surrounded by an endoneurium, and groups of endoneuria are encased by a perineurium to form a fascicle. The fascicles and blood vessels are in turn surrounded by the epineurium to complete the nerve. Below the schematic is a representative IHC slice showing localization of Na_V_1.7 in the plasma membrane of the nerve fibers. (b) Quantification of the extent of Na_V_1.7 expression (% of toal nerve fibers?) in human peripheral nerves from males (n = 3) and females (n = 3). (c, d) Representative IHC slices showing expression of Na_V_1.7 in nerves from the torso and limbs of male and female cadavers, respectively. (e, f) Quantification of Na_V_1.7 staining in panels c and d, respectively. (g) Comparison of Na_V_1.7 expression in motor and sensory nerves of humans (males + females). (h) Comparison of Na_V_1.7 expression in upper- and lower-body mixed nerves. (i) IHC staining of Na_V_1.7 in mouse brachial plexus. (j) Quantification of Na_V_1.7 expression in different mouse organs. (k) IHC staining of Na_V_1.7 in NHP vagus and recurrent nerves. (l) Quantification of Na_V_1.7 expression in different NHP organs. (m) Repurposing approach to engineer natural peptides for use in surgeries. Scheme showing a mixture of peptides, isolation, engineering, formulation and the much-needed accessing of these peptides to the peripheral nervous system.

The majority of nerves showed high or very high NaV1.7 expression. Especially very high expression was noted in branches of the trigeminal nerve, accessory nerve, and some branches of the genitofemoral nerves, and high expression was observed in the vagus and recurrent laryngeal nerves (Figure 1c-f). We found no difference in Na_V_1.7 expression between the motor and sensory nerves (Figure 1e). The medium-expression nerves were mostly from the extremities, including the radial, median, and tibial nerves. Despite some variability, in most cases Na_V_1.7 expression was high in nerves compared to other tissues, which were accompanied by H&E staining showing circular dots or transversal tracks (Figure S1b). Some minor expression was noted in the thyroid gland and myocardium, but other striated and smooth muscle tissues had almost no expression. Furthermore, we did not observe any difference in expression between torso and limb nerves or left or right limbs (Figures 1f-h). We also assessed Na_V_1.7 expression in mouse nerves and other organs (Figure 1i and 1j) using IHC. Some Na_V_1.7 expression was observed in the liver (20.4 ± 7%), the expression was lower when compared to nerves, though not statistically significant (26.1 ± 11.6%, p > 0.05). Minimal expression was observed on the other analyzed organs (muscle = 14 ± 8, heart = 9 ± 4, spleen = 15 ± 11, kidney = 17 ± 7, and brain = 10 ± 6).

To confirm cross-species conservation of expression, we conducted comparative studies in NHPs and evaluated differences in Na_V_1.7 expression across various nerves and organs. (Supplementary Table S1). This included IHC staining of the sciatic, vagus, and recurrent laryngeal nerves, as well as the muscle, heart, spleen, liver, kidney, brain, and dorsal root ganglia (DRG) (Figures 1k, 1l). IHC staining showed Na_V_1.7 signals on the DRG and the kidneys. The specificity of the anti-Na_V_1.7 antibody in both species was confirmed by the absence of staining on IgG control slides (Figure S2).

### Engineering a library of fluorescent Tsp1a conjugates

To prepare tracers, we modified Tsp1a synthetic peptides with fluorophores, including IR800, Janelia669, Bodipy665, DY684, and CY7.5 (Figure 2a), generating fluorescent, nerve-targeted tracers based on Tsp1a. Specifically, we chose to modify Tsp1a-K4 and Tsp1a-Pra0 peptides by nucleophilic substitution and by dipolar cycloaddition, respectively; analogous to our previous work ^51,56^. Tsp1a-K4 and Tsp1a-Pra0 peptides were modified with IR800, Janelia669, Bodipy665, DY684, and CY7.5 using either N-hydroxy succinimide (NHS) ester or an azide moiety for the corresponding conjugations (Figure 2b, 2c). These fluorophores were selected due to their different sizes, charge, relatively high stability, and optical relevance for use in the operating room. For Tsp1a-K4, the chemical transformations were performed under basic conditions in a mixture of water and acetonitrile, yielding the corresponding conjugates Tsp1a-IR800_K4_, Tsp1a-JA669_K4_, Tsp1a-BO665_K4_, Tsp1a-DY684_K4_ and Tsp1a-CY7.5_K4_ in 42%, 40%, 32%, 22%. These efforts led to 40% isolated yields with 95% purity for all engineered tracers. For Tsp1a-Pra0, the chemical transformations were performed under basic conditions (pH 9–10) in a combined solution of Tris-buffer, ascorbic acid, and copper sulfate. This yielded the corresponding probes Tsp1a-IR800_P_, Tsp1a-JA669_P_, Tsp1a-BO665_P_, Tsp1a-DY684_P,_ and Tsp1a-CY7.5_P_. These efforts led to 55%, 46%, 40%, 28% and 44% isolated yields, respectively, with 95% purity for all conjugates.

**Figure 2.**
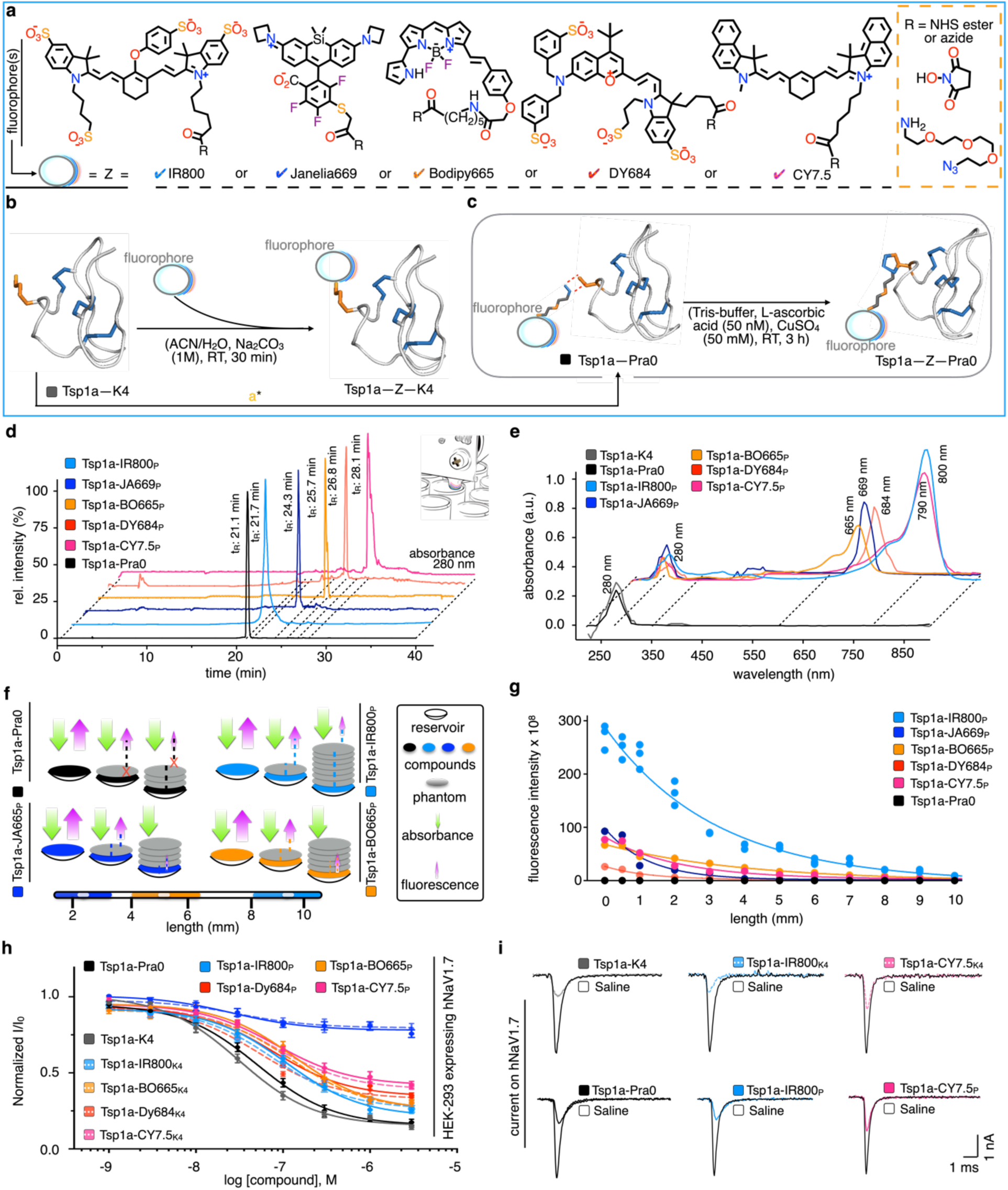
Synthesis of a library of Na_V_1.7-targeted fluorescent tracers. (a) Structures of clinically-relevant fluorophores used in this study: IR800, Janelia669, Bodipy665, Dy684 and Cy7.5. (b, c) Reaction schemes for conjugation of red and near-infrared dyes to Tsp1a via either (b) NHS ester coupling to the sidechain of residue K4 or (c) dipolar cycloaddition to a non-native propylglcyine residue (Pra0) added to the N-terminus of Tsp1a. (d) RP-HPLC chromatograms showing retention times of of Tsp1a-Pra0, Tsp1a-IR800_P_, Tsp1a-JA669_P_, Tsp1a-BO665_P_, Tsp1a-DY684_P_ and Tsp1a-CY7.5_P_ peptides. Peptide absorbance was monitored at 280 nm. (e) Absorbance spectra of 0.2 μM Tsp1a-Pra0, Tsp1a-IR800_P_, Tsp1a-JA669_P_, Tsp1a-BO665_P_, Tsp1a-DY684_P_ and Tsp1a-CY7.5_P_ peptides showing expected absorption maxima for each of the attached fluorophores. (f) Fluorescence intensity obtained from Tsp1a-Pra0 peptides in the presence of phantoms with increasing thickness in mm (g) Quantification of fluorescence penetration of Tsp1a peptides. Tsp1a-IR800_P_ had significantly better penetrance than any other tracer. (h) Concentration-response curves for inhibition of hNa_V_1.7 by fluorescently-labeled Tsp1a-Pra0 peptides. (i) Representative hNa_V_1.7 currents in the presence or absence of Tsp1a peptides.

For Tsp1a-K4 conjugates, the retention times (r_t_) shifted from 21.1 min for Tsp1a-K4 to 21.6 min, 24.3 min, 25.8 min, 24.2 min and 28.0 min for Tsp1a-IR800_K4_, Tsp1a-JA669_K4_, Tsp1a-BO665_K4_, Tsp1a-DY684_K4_ and Tsp1a-CY7.5_K4_, respectively (Figure S4a, b, e). For Tsp1a-JA669_K4_, Tsp1a-BO665_K4_, Tsp1a-DY684_K4_ conjugates, the major impurities were the partially reduced peptide products, 5-9% (r_t_ 26.3 min, 27.8 min and 28.8 min, respectively), which were also present in the starting materials (r_t_ 21 min, 92% and r_t_ 21.2 min, 8% for Tsp1a-K4 and reduced Tsp1a-K4, respectively). For Tsp1a-Pra0 fluorescent conjugates, the retention times (r_t_) shifted from 21.1 min for Tsp1a-Pra0 to 21.7 min, 24.3 min, 25.7 min, 26.8 min and 28.1 min for Tsp1a-IR800_P_, Tsp1a-JA669_P_, Tsp1a-BO665_P_, Tsp1a-DY684_P_ and Tsp1a-CY7.5_P_, respectively (Figure 2d and Figure S3). Standard LC-MS experiments were performed for all Tsp1a tracers and digestions for Tsp1a-K4 peptides, (Figures S4, S5).

### Photophysical features of fluorescently labeled Tsp1a-K4 and Tsp1a-Pra0 conjugates

We expected a dual peptide/dye absorbance peak for all of the fluorescently-labeled conjugates. The absorption and emission spectra of Tsp1a-K4 and Tsp1a-Pra0 probes were also determined. For absorption, the characteristic photo-physical red and near-infrared ex_max_ = 665 nm, ex_max_ = 669 nm, ex_max_ = 684 nm, ex_max_ = 780 nm and ex_max_ = 800 nm signals for the fluorophores families of Bodipy665, Janelia669, DY684, CY7.5 and IR800 were observed. These values were used as criteria to confirm dye conjugation in all engineered peptides. These results, coupled with the observation of peptide-absorbance peaks at around 250–280 nm were used to further confirm conjugation of Tsp1a to the fluorophore. We expected all fluorescently labeled conjugates to have a dual peptide/dye absorbance peak at 280 nm and absorbance signals (Soret bands) at the corresponding red and near-infrared region. This was observed for all fluorescent engineered conjugates of Tsp1a-K4 and Tsp1a-Pra0 (Figures 2e, S6a).

### Depth penetration testing with phantoms, tissue-like fluorescence infiltration

For our penetration experiments, we designed protocols that could better highlight the photophysical properties of our Tsp1a tracers while concomitantly considering the routinely-used reputed fluorophores and clinically-relevant tools to develop our nerve imaging agent and help choose a lead candidate. A 1 mm phantom was prepared and overlayed on top of a reservoir containing the corresponding engineered fluorescent tracer (50 µL of 1 nmol, 10 µM of Tsp1a-tracer in 100 µL PBS). The thickness of the phantom(s) was measured, and we assayed the penetration power of each fluorescent Tsp1a tracer (Figures 2f, g, S6b). We found out that >9 phantoms (9 mm thickness) did not prevent the emission of Tsp1a-IR800_P_ tracers which ranked with the highest penetration power. 7 phantoms were required to obstruct the emission of Tsp1a-BO665_P_ and Tsp1a-CY7.5_P_, respectively. 2 and 3 phantoms were used to obstruct the emission of Tsp1a-DY684_P_ and Tsp1a-JA669_P_, respectively, (Figure 2g).

### Inhibition of human Na_V_1.7 by Tsp1a and fluorescently-labeled conjugates

We examined the inhibitory effect of Tsp1a-K4 and Tsp1a-Pra0 peptides and derivatives on a panel of human Na_V_ subtypes by using automated whole-cell patch-clamp electrophysiology. Both Tsp1a-K4 and Tsp1a-Pra0 retained the inhibitory effect on human Na_V_1.7 (IC_50_ = 34–50 nM), and their potencies were only marginally reduced by the addition of NIR fluorophores to the core Tsp1a peptide (IC_50_ = 50–140 nM) (Figures 2h, 2i, S7). Since the fluorescent Tsp1a Pra0 peptides are nanomolar inhibitors of Na_V_1.7, they should be suitable for targeting peripheral nerves in mice, NHPs and humans. Tsp1a-IR800_P_ also showed no affinity for other sodium channels (Figure S8). Table 1S shows the most important features of Tsp1a tracers.

### Histological validation of Na_V_1.7 expression

IHC staining showed high Na_V_1.7 expression and nerve track continuity in mouse nerve tissue and elongated nerve tracks in H&E adjacent samples. We found no expression of Na_V_1.7 in mouse muscle, heart, spleen, or brain. In contrast, the signal was positive in mouse kidney and liver (Figure 3a). Thus, the IHC data validate Na_V_1.7 as a biomarker of peripheral nerves in mice. Figure 3b shows an overview of the *in vivo* approach used to observe repeated accumulations of Tsp1a tracers in mouse thicker nerves.

**Figure 3.**
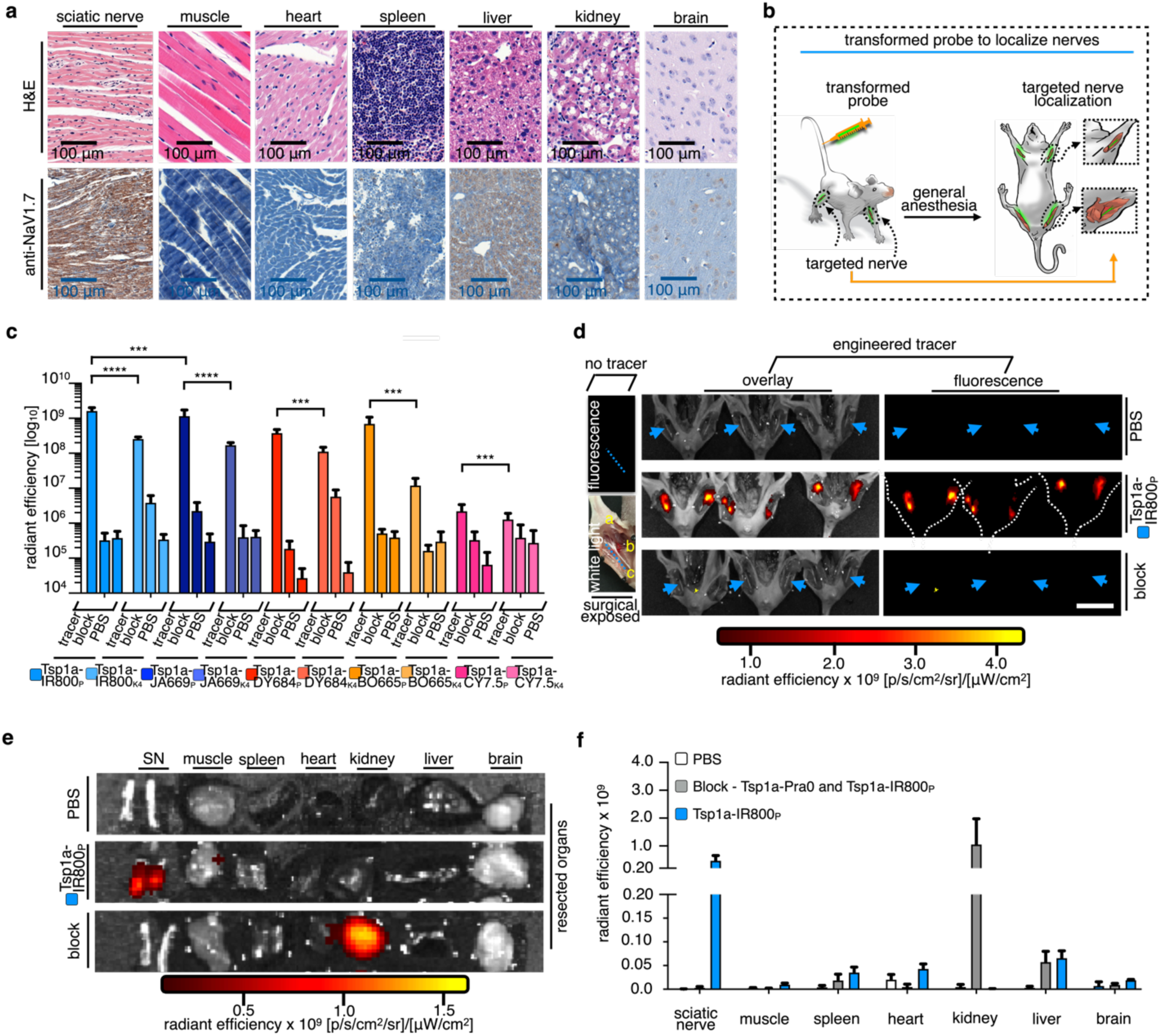
Na_V_1.7 expression in mice and pharmacokinetic evaluation of fluorescently-labeled Tsp1a peptides. (a) Hematoxylin and oesin (H&E) and anti-Na_V_1.7 IHC staining of sciatic nerves and resected organs from mice. (b) Schematic representation of Tsp1a tracer injection and anticipated map for the imaging of peripheral nerves. (c) Quantification of fluorescence intensity of exposed peripheral nerves from mice injected with vehicle, Tsp1a tracer (i.e., Tsp1a-Pra0 and Tsp1a-K4 conjugates, 10 µM in 100 µL PBS), or block formulation containing both tracer and a 20-fold excess of ‘cold’ unconjugated Tsp1a). (d) White light and fluorescence images of sciatic nerve with no Tsp1a tracer (left) and epifluorescence images (right) of animals injected with 100 µL PBS vehicle (top), Tsp1a-IR800_P_ (1 nmol; 10 µM Tsp1a-IR800_P_ in 100 µL PBS) (middle) or a Tsp1a-IR800_P_/Tsp1a-Pra0 block formulation (Tsp1a-IR800_P_, 10 µM, 1 nmol and Tsp1a-Pra0, 204 µM, 21 nmol in 100 µL PBS) (bottom). Images of exposed peripheral nerves of mice were taken 30 min after tail vein injection. (e) Epifluorescence images of resected right and left peripheral nerves and organs from mice injected with PBS, Tsp1a-IR800_P_ or a block formulation. High fluorescence intensity (due to fluorophore accumulation) was only observed in sciatic nerves from mice injected with Tsp1a-IR800_P_ alone. No fluorescence was observed after 30 min for mice injected with vehicle or block formulation, except for some accumulation in the kidney of mice injected with block formulation (an expected consequence of renal clearance of the tracer). (f) Quantification of radiant efficiency in sciatic nerve and organs of mice injected with PBS, Tsp1a-IR800_P_ or block formulation. Statistics were calculated using

### Pharmacokinetics of the engineered fluorescent Tsp1a peptides in mice

In our microscale imaging experiments, we observed rapid and selective accumulation of Tsp1a tracers in mice (Figure 3c). Mice were injected via the tail vein with a fluorescent Tsp1a tracer alone (1 nmol, 10 µM in 100 µL of PBS), Tsp1a tracer in combination with an excess of the native Tsp1a peptide (21 nmol, 204 µM in 100 µL of PBS) (block), or with vehicle (PBS). Mice were sacrificed 30 min after injection. The thickest peripheral nerves were chosen for surgical exposure and observation of tracer accumulation. The right and left sciatic nerves (RSN and LSN) were surgically exposed (Figures 3d, S9), and epifluorescence imaging was performed using an IVIS Spectrum *in vivo* imaging system (excitation 780/20-800/20 nm; emission 800–840 nm, see SI for full list of excitations and emission of the other engineered conjugates). When the Tsp1a-IR800_P_ imaging agent was injected in mice, the sciatic nerves were visible, and a high fluorescence signal was observed when compared to the block group (radiant efficiency: 1.62 ± 0.39 x 10^9^ and 0.0003 ± 0.0002 x 10^9^ for sciatic nerves, respectively; Student’s unpaired *t*-test, ****P < 0.0001). The PBS control group had a radiant efficiency of 0.0004 ± 0.0002 x 10^9^ for sciatic nerves. For the other Tsp1a tracers, similar observations were noted when generating contrast between surrounding tissue and the sciatic nerves. The tracers accumulated on sciatic nerves within 30 min of intravenous injection (Figures S9, S10). The conjugates with Cy7.5 fluorophore did not generate strong contrast, which may be due to lack of retention on nerves and overall charge of the construct.

On the ex-vivo and biodistribution, the differences in radiant efficiency were equally pronounced (Figures 3e, f, S10). The sciatic nerves are visible in mice receiving the imaging agent only, showing an average radiant efficiency of 0.5 ± 0.2 x 10^9^. In contrast, mice receiving the imaging agent with the unmodified Tsp1a peptide (21 nmol, 204 µM in 100 µL of PBS) had a statistically significant 149-fold reduction in radiant efficiency to 0.003 ± 0.003 x 10^9^ (Student’s unpaired *t*-test, ***P < 0.0001).

### Measuring the intraoperative window of Tsp1a-IR800_P_

The *in vivo* circulation of Tsp1a-IR800_P_ was observed and measured in a blood half-life experiment using an *in vivo* imaging system (IVIS). After retro-orbital blood withdrawal from animals injected with Tsp1a-IR800_P_, the blood half-life was measured to be approximately 3 min (Figure 4a); the tracer washed out slowly after 2 h. Furthermore, we traced the accumulation of Tsp1a-IR800_P_ in the kidneys *in vivo* using the IVIS (Figure 4b) and tracked the circulation of the Tsp1a-IR800_P_ and its metabolites via HPLC and LC-MS analysis of blood samples. Our HPLC results showed a detectable signal at approximately 6.1 min. However, this signal was not present in purified Tsp1a-IR800_P_ or the blood of non-injected animals (Figure 4c; 280 nm and 800 nm wavelengths) (Figure S5a). Our LC-MS results showed a large ion signal at 1563, corresponding to [M+3]^3+^ from the fluorescent tracer, and m/z signals at 689 and 781, corresponding to time points, 5 min and 10 min, respectively (Figure 4d).

**Figure 4.**
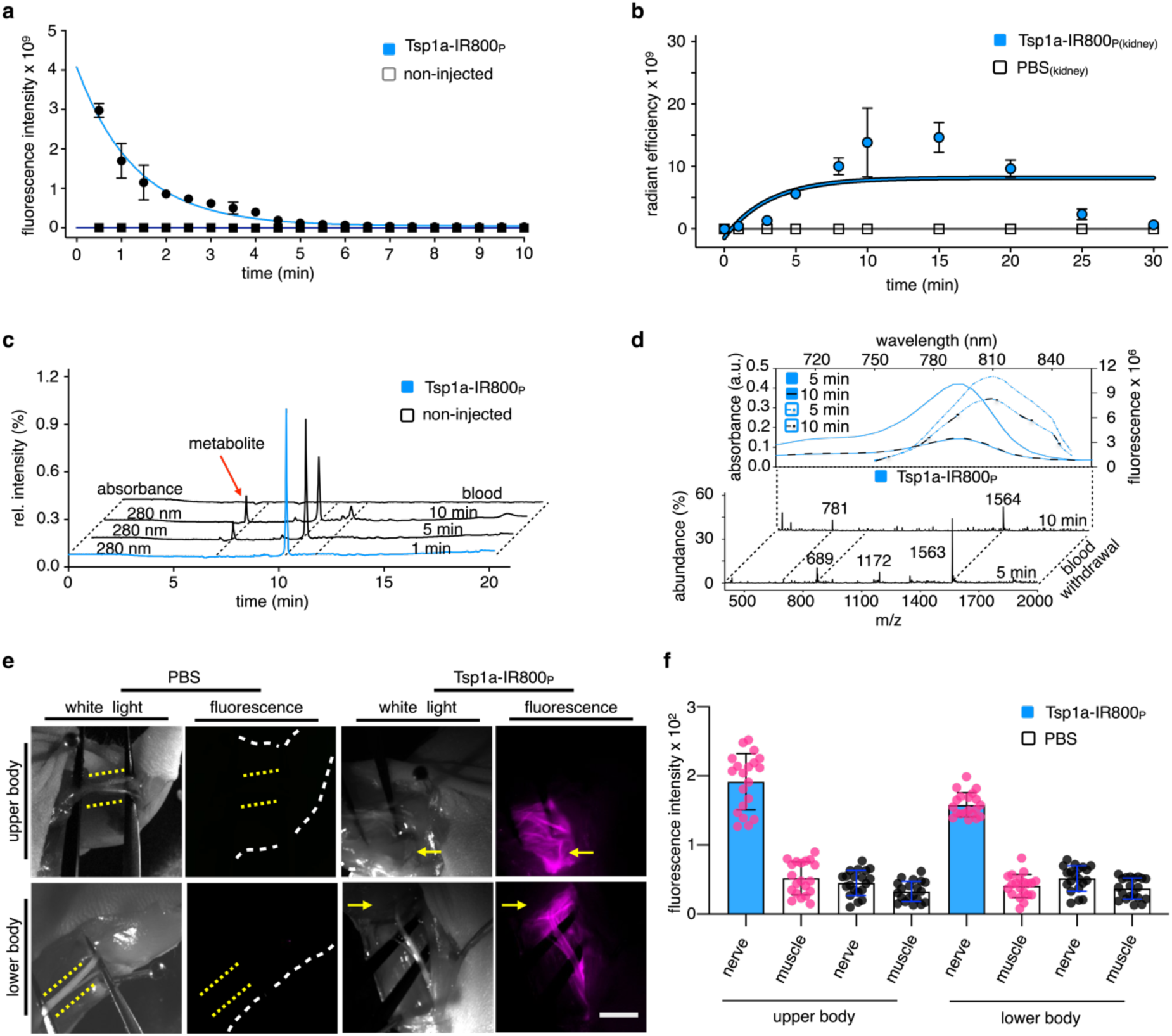
Imaging of sciatic nerves in mice injected with Tsp1a-IR800_P_ . (a) Half-life of Tsp1a-IR800_P_ in mice (injected dose: 1 nmol; 10 µM Tsp1a-IR800_P_ in 100 µL PBS) based on quantification of fluorescence intensity. (b) Fluorescence tracking of Tsp1a-IR800_P_ (1 nmol, 10 µM of Tsp1a-IR800_P_ in 100 µL PBS) in kidney. (c) Metabolism of Tsp1a-IR800_P_ in mouse serum (injected dose: 1 nmol; 10 µM Tsp1a-IR800_P_ in 100 µL PBS) tracked using RP-HPLC. A minor metabolite appeared at shorter retention time 5 min after tracer injection. (d) Metabolism of Tsp1a-IR800_P_ in mouse serum (1 nmol; 10 µM Tsp1a-IR800_P_ in 100 µL PBS) tracked using LC-MS. (e) Stereoscopic images (Zeiss Lumar stereoscope) of exposed sciatic nerves from mice that were injected with Tsp1a-IR800_P_ (1 nmol; 10 µM Tsp1a-IR800_P_ in 100 µL PBS) or PBS under white light and fluorescent excitation. Images were taken in brightfield and with 750 nm laser excitation (exposure time 250 ms). (f) Quantification of fluorescence images in panel e (n = 3 per group). Statistics were calculated using a nonparametric Student’s *t*-test. \**P* < 0.01. Error bars are standard deviations.

### Surgical microscope imaging of Tsp1a-IR800_P_

Mice were injected with Tsp1a-IR800_P_ (1 nmol, 10 µM in 100 µL of PBS) or 100 µL of PBS and immediately put under anesthesia with isoflurane. After 5 min, mouse nerves were exposed for imaging using a Lumar surgical fluorescence stereoscope (SteREO Lumar.V12, Zeiss, Jena, Germany). (Figures 4e, 4f). Figure 4e shows images of the brachial plexus (top row) and sciatic nerves (bottom row) obtained under white light imaging conditions (first and third columns) and under fluorescence imaging conditions (second and fourth columns) for mice injected with Tsp1a-IR800_P_ or PBS. Unlike the IVIS Spectrum, the Lumar Fluorescence imaging system does not support autofluorescence suppression. We detected strong specific fluorescence in the nerves of animals injected with Tsp1a-IR800_P_ (fluorescence intensity: 1.8 ± 1.7 x 10^2^ and 2.1 ± 0.5 x 10^2^ for the brachial plexus and sciatic nerves, respectively), which was significantly lower for PBS-injected mice (0.7 ± 0.2 x 10^2^ and 0.4 ± 0.2 x 10^2^, for brachial plexus and sciatic nerves, respectively, Student’s test, P < 0.05, Figure 5f).

**Figure 5.**
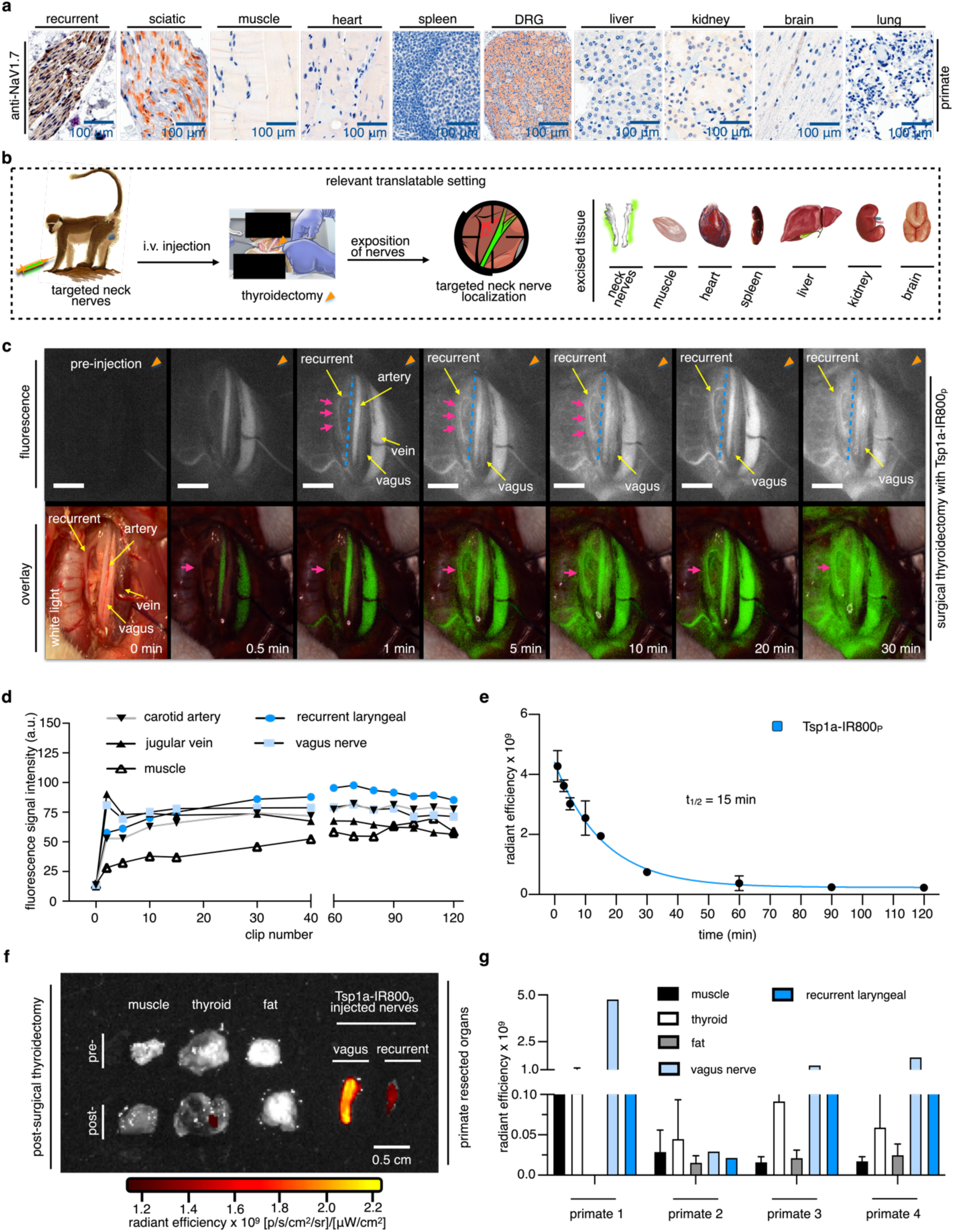
Performance of Tsp1a-IR800_P_ in a clinically translatable setting. (a) Anti-Na_V_1.7 immunohistochemical staining showing that Na_V_1.7 is highly expressed in the peripheral nerves (recurrent laryngeal, sciatic, and DRG) but not resected organs from NHPs. (b) Schematic representation of workflow used for imaging of peripheral nerves with Tsp1a tracer during thyroidectomy (orange triangle) in NHPs. (c) Quest images of a thyroidectomy with primates, fluorescence and overlay images of primates injected with Tsp1a-IR800_P_ (250 µg/kg in 5 mL PBS) which was administered intravenously (5 mL in ∼60 s), images of exposed left peripheral nerves in primates showing the recurrent laryngeal nerve lighting up (pink arrows), followed by demarcation of the vagus nerve over 30 min. Peripheral nerves are labeled (yellow arrows). (d) Quantification of fluorescence intensity from video fragments of exposed recurrent laryngeal nerve, vagus nerve, and surrounding tissues following injection of Tsp1a-IR800_P_ (250 µg kg^−1^, in 5 mL of PBS) in NHPs. (e) Half-life of Tsp1a-IR800_P_ (250 µg kg^−1^, in 5 mL of PBS) in NHPs determined from fluorescence of blood samples. (f) Epifluorescence images of resected peripheral nerves and surrounding tissues from NHPs following injection of Tsp1a-IR800_P_ (250 µg/kg in 5 mL PBS). High fluorescence intensities (due to Tsp1a-IR800_P_ accumulation) were observed in primate peripheral nerves from the thyroidectomy. (g) IVIS fluorescence quantification of resected tissues from primates injected with Tsp1a-IR800_P_ (250 µg/kg in 5 mL PBS). Statistics were calculated using a non-parametric Student’s *t*-test. *P < 0.05; **P < 0.01; ***P < 0.001; ****P < 0.0001.

### Performance of Tsp1a-IR800_P_ in a clinically translatable setting

Our IHC studies revealed high expression of Na_V_1.7 in nerves from NHPs, including the recurrent laryngeal, vagus, and sciatic nerves (Figure 5a). Accompanied by H&E staining (Figure S12), we also noted expression of Na_V_1.7 in a few other organs, namely the medulla and cortex of the brain, and the kidney (Figures 1l and 5a). These results validated Na_V_1.7 as a biomarker of peripheral nerves in primate models.

We conducted an experiment simulating human thyroidectomy in NHPs, a surgery in which accidental injury of the recurrent laryngeal nerve (RLN), a branch of the vagus nerve that innervates all of the intrinsic muscles of the larynx except the cricothyroid muscles, is common. Indeed, the incidence of RLN injury (temporary and permanent) after thyroid and parathyroid surgery was recently estimated to be exceptionally high reaching almost 14%, and depended on the volume and experience of the surgeon.^57,58^

We used Tsp1a-IR800_P_ as an imaging tool to guide the surgery (Figure 5b). During the thyroidectomy in NHPs, we observed rapid and selective accumulation of Tsp1a-IR800_P_ in the RLN within 1 min, followed by slower accumulation in the vagus nerve at ∼5 min (Figure 5c, S12). (Figure 5c, S12).

The average fluorescence intensity over a 40-min period was measured for various anatomical structures: muscle (25.3 ± 1.2), carotid artery (60.4 ± 1.4), jugular vein (79.7 ± 1.1), RLN (85.4 ± 1.3), and vagus nerve (88.7 ± 1.1), with statistically significant difference between surrounding muscles and nerves (P < 0.001, Student’s unpaired *t*-test, Figure 5d). Simultaneously, blood withdrawal facilitated the determination of the tracer half-life in primates (t_1/2_ = 15 min), revealing a moderately fast clearance (Figure 5e). For *ex vivo* validation, we examine the resected primate nerve tissue and organs. The differences were notable in the peripheral nerves compared to the surrounding tissue (Figures 5f, S13), with the exception of primate 2, where tracer administration was not stellar due to technical difficulties encountered during the injection (Figure 5g). We have not observed significantly higher fluorescent signals in other organs, including muscle, fat and thyroid (Figure 5g).

### The safety profile of Tsp1a-IR800_P_ in mice and NHPs

We monitored critical parameters and relevant vital signs of mice before injection, immediately after injection, and 5 min after injection of Tsp1a-IR800_P_ (1 nmol, 10 µM in 100 µL of PBS) or 100 µL of PBS with a rodent surgical monitor (Scintica, Instrumentations, Houston). These included general physical signs, as well as oxygen levels, heart rates, and body temperatures (Figure 6a-d). During NHP surgeries, no abnormal changes and behavior were observed when the animals were injected with Tsp1a-IR800_P_. Figures 6e-g show no toxicity confirming the safety of Tsp1a-IR800_P_.

**Figure 6.**
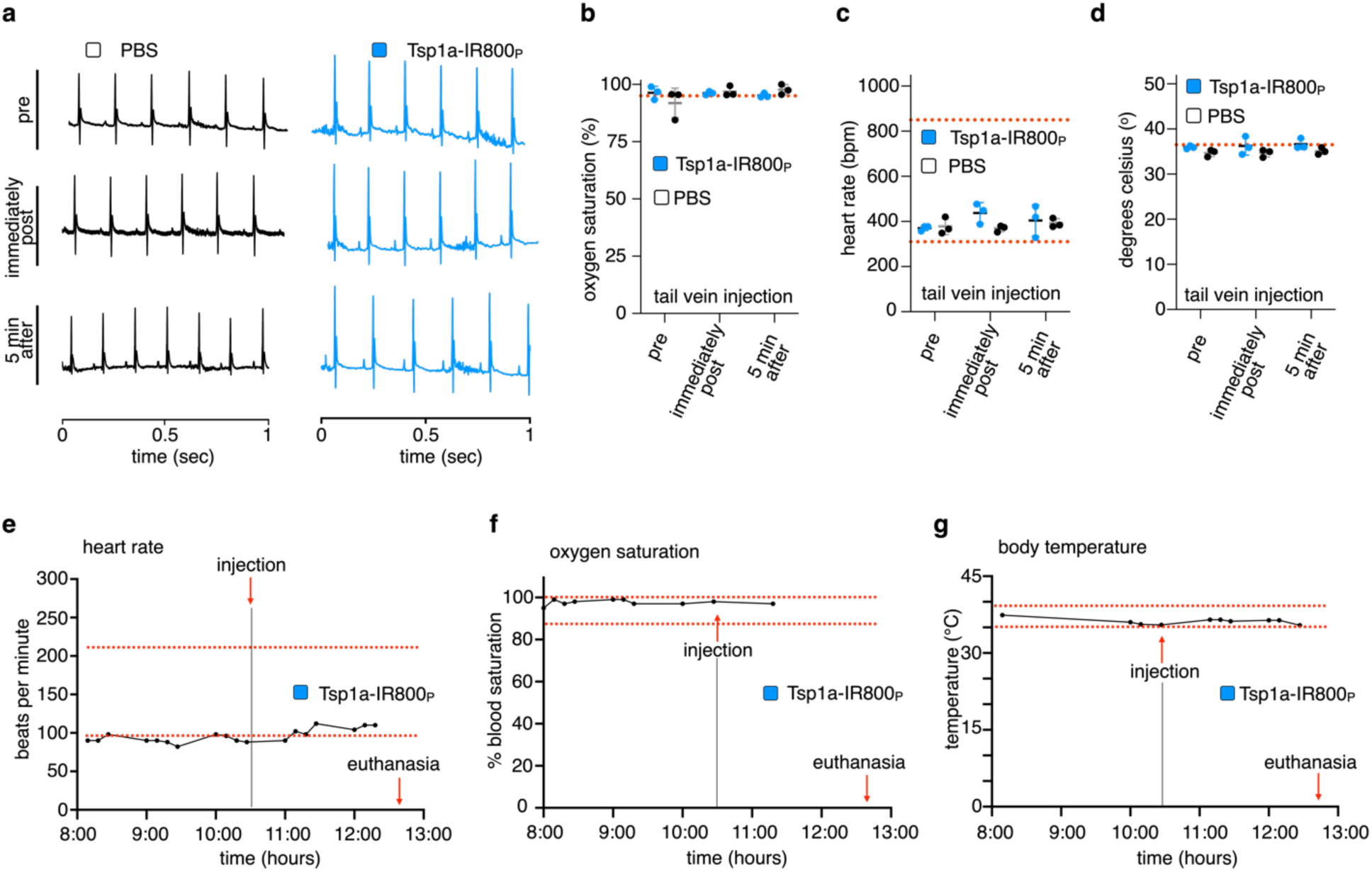
Pharmacodynamics of Tsp1a-IR800_P_ in mice and NHPs. (a) Electrocardiograms from mice injected with Tsp1a-IR800_P_ (20 µg/kg in 100 µL PBS) showing normal cardiac electrical activity before and after injection of the tracer. Injection of Tsp1a-IR800_P_ into mice also had no impact on (b) oxygen saturation, (c) heart rate or (c) body temperature. No abnormal behavior or other overt signs of toxicity were observed after intravenous administration of Tsp1a-IR800_P_ in mice. Monitoring of (e) oxygen saturation, (f) heart rate and (g) during thyroidectomy in NHPs showed no impact from Tsp1a-IR800_P_ adminstration. No abnormal behavior or other overt signs of toxicity were observed following intravenous administration of Tsp1a-IR800_P_ and during the surgical procedure.

## DISCUSSION

IND provides a challenge even for highly skilled surgeons. Physicians often encounter obstacles due to changes induced by tumor progression or invasion, deformed anatomy, previous radiation therapy or surgery. These factors result in scarred tissue textures or atypical planes, leading surgeons to rely heavily on visual feedback, palpation, experience, and anatomical landmarks, which can yield unreliable assessments of nerve location. A potential solution to this problem is to take advantage of bioactive peptides that selectivity target neuronal receptors to allow fluorescence-guided intraoperative imaging of buried nerve tissues.

Currently, several fluorescent cancer-targeted agents are in the clinical translation pipeline with emission maxima of 750–800 nm. Leveraging this idea, we believe nerve-targeted contrast agents can utilize similar equipment and fluorescent properties for use in nerve localization for easy deployment to modern operating theaters. In this study, we developed nerve-targeted agents based on the conjugation of fluorophores to Tsp1a, a serum-stable disulfide-rich venom peptide that targets Na_V_1.7 with exquisite potency and selectivity. Importantly, Tsp1a does not target closely related Na_V_ channels that have critical physiological functions and different anatomical locations, such as Na_V_1.4 in muscle and Na_V_1.5 in the heart. Moreover, the peptide’s nanomolar affinity for Na_V_1.7 should enable OS to generate sufficient contrast with micro-doses of Tsp1a tracer, in contrast to the much higher doses that would be required for therapeutic targeting of Na_V_1.7, thereby further ameliorating the potential for adverse side-effects and inadvertent targeting of non-neuronal tissue.

We underpinned this work by exploring the distribution of Na_V_1.7 in cadaveric human tissue to provide validation of biomarker presence, and we confirmed the anticipated high abundance of Na_V_1.7 in human peripheral nerves in both males and females. Moreover, we demonstrated high levels of Na_V_1.7 expression in nerves from mice, non-human primates (grivets) and humans, indicating that the channel is well conserved across species.

Encouragingly, conjugation of a wide range of fluorophores to Tsp1a did not significantly impact the peptide’s affinity for Na_V_1.7 or its selectivity for Na_V_1.7 over other Na_V_ channel subtypes. The fluorescent Tsp1a tracers generated high fluorescence signals (2 x 10^9^ radiant efficiency) to visualize nerves, which were corroborated with stereoscopic imaging *in vivo*. Following i.v. administration, the Tsp1a-IR800_P_ tracer, which had the highest penetrance, accumulated in the upper (brachial plexus) and lower (sciatic) extremity nerves. Strikingly, fluorescence delineation of upper- and lower-body extremity nerves was observed with current standard-of-care clinical cameras (Lumar stereoscope and Quest camera). Peripheral nerves from mice injected with Tsp1a-IR800_P_ produced bright, high-contrast fluorescence to guide surgery, rendering an immediate semi-quantifiable readout. These results are very encouraging because, despite clinical cameras not allowing image deconvolution (and therefore autofluorescence correction), Tsp1a-IR800_P_ accumulation was clearly resolvable and confirmed our previous nerve visualization studies. Other Na_V_1.7-targeted tracers such as Tsp1a-JA669_P_ also performed well *in vivo*, showing good delineation of nerves.

Nerve visualization with Tsp1a tracers was also used to successfully guide thyroidectomy surgery in NHPs. Following i.v. administration of Tsp1a-IR800_P_, fluorescence signals were obtained in exposed laryngeal recurrent and vagus nerves which provided clarity in the normal surgical dissection plane and enabled these nerves to be differentiated from large vessels and adjacent tissues. Importantly, Tsp1a-IR800_P_ caused no toxicity in either mice or primates. Mice were unaffected by doses of 5–45 µg/kg Tsp1a-IR800_P_ and primates tolerated a dose of 250 µg/kg Tsp1a-IR800_P_ with no adverse physical signs observed during the surgical intervention.

The Tsp1a tracers have limitations in terms of competing with fluorescence from other structures including large vessels and accumulation on adjacent tissue in NHPs. These hurdles are intrinsic to new imaging approaches which require optimization to tailor the agent for different procedures and surgical scenarios. Nevertheless, we believe that Tsp1a-IR800_P_ should stimulate a paradigm shift in intraoperative imaging of peripheral nerves to avoid IND. It conveniently enables visualization of nerves within 20 min of injection, which could provide an immediate *in vivo* readout for urgent surgical applications in the OR.

In summary, we engineered a library of Na_V_1.7-targeted fluorescent tracers with desirable physiochemical properties, an ability to accumulate in nerves, and the capacity for nerve visualization during surgery. The most penetrant tracer, Tsp1a-IR800_P_, enabled clear delineation of the recurrent laryngeal and vagus nerves during thyroidectomy in NHPs. Administration of Tsp1a-IR800_P_ could be implemented in the OR for routine nerve localization for patients undergoing surgery to minimize the risk of iatrogenic nerve damage.

### Statistical analyses

Statistical analyses were performed using GraphPad Prism 8. Unless otherwise stated, data points represent mean values, and error bars represent standard deviations of biological replicates. P values were calculated using a Student’s unpaired t-test or Chi-square.

## Acknowledgments

The authors thank the Imaging and Radiation Sciences Program and the MSK Molecularly Targeted Intraoperative Imaging Fund, the Small Animal Imaging Core (P. Zanzonico, V. Longo), the Radiochemistry and Molecular Imaging Probes Core (S. Lyashchenko), and the Molecular Cytology Core at Memorial Sloan Kettering Cancer Center for support. The authors also thank Dr. Snehal Patel, Dr. Ian Ganly, Dr. Alvin Goh, Dr. Elizabeth Jewel, Dr. Daniela Molena and Dr. Lucas M. Nogueira for help with nerve dissection. We also thank the lab techs who were very instrumental in multiple aspects of this study Alexa Michel, Navjot Guru, and others. The funding sources were not involved in the study design, data collection and analysis, writing of the report, or the decision to submit this article for publication. All authors carefully read and edited the manuscript.

## Competing Interests

T.R. and J.S.L are shareholders of Summit Biomedical Imaging, LLC. T.R. is a paid consultant for Theragnostics, Inc. T.R. is now an employee of and a shareholder of Evergreen Theragnostics. All other authors have no conflict to declare. This arrangement has been reviewed and approved by MSK in accordance with its conflict-of-interest policies. P.D.S.M., J.G, J.S.L., T.R., Y.J., and G.F.K. are co-inventors on a US patent describing the Tsp1a peptide.

## Funding

This work was supported in part by the National Institutes of Health grants R01 EB029769, R00 GM145587, R35CA232130, and P30 CA008748, grant agreement No 796672, the Australian National Health and Medical Research Council (Project Grant APP1080405, Program Grant APP1072113 to G.F.K. and the Australian Research Council (Centre of Excellence grant CE200100012 to G.F.K). We are extremely grateful for funding from Mr. William H. Goodwin and Mrs. Alice Goodwin and the Commonwealth Foundation for Cancer Research and The Center for Experimental Therapeutics of MSKCC (T.R. & N.P.) and Technology Development Fund (N.P.) of MSKCC is acknowledged, that facilitated conducting primate studies described here in the manuscript. No NIH funds were used for conducting primate studies.

## Author Contributions

J.G., T.R., and N.P. conceived the study and designed the experiments. J.G., P.D.S.F., D.A., R.A., C.Y.C, and T.V. and D.S.J. carried out the experiments and collected the data. G.F.K., C.Y.C., C.I.S., and Y.J. contributed new reagents/analytical tools. J.G., D.A., P.D.S.M., C.Y.C., G.F.K., J.S.L., A.C.G., D.M., J.E., S.G.P, T.R. and N.P interpreted the data. J.G., P.D.S.F., D.A., N.P., and T.R. wrote Institutional Research Board protocols. P.D.S.F., D.A., J.G., and N.P. led NHP’s experiments. D.A., P.D.S.F., performed human nerve dissections and thyroidectomies. J.G., D.A., P.D.S.F., T.R., and N.P. wrote and edited the manuscript. All authors carefully read and edited the manuscript.

