## Supplementary Information for "Na_V_1.7 targeted fluorescence imaging agents for nerve identification during intraoperative procedures"

### RESULTS

#### $Na_V$1.7 expression is conserved across mice, humans and NHPs

$Na_V$1.7 is a voltage-gated sodium channel that is highly expressed in peripheral sensory neurons and mixed nerves. Nerve bundles are intricate structures made of a network of nerve fibers (axons) encased by Schwann cells and surrounding by a protective endoneurium. Groups of endoneuria are encased by a perineurium to form a fascicle. A bundle of fascicules and blood vessels are then encased by a layer of connective tissue called epineurium to form a mesoscopic nerve structure. $Na_V$1.7 is expressed throughout the nerve fibers within each of the fascicles and it is critical for the transmission of pain signals to the spinal cord (Figure 1a).

#### Histological validation of channel $Na_V$1.7 and experimental design

In previous work, we validated the expression of $Na_V$1.7 in the peripheral nerves of mice. In the current study, using H&E and anti-$Na_V$1.7 staining of nerve tissue, resembling the histological structures obtained previously with circular structures, identified as nerve bundles when slicing transversally and when slicing the nerve was longitudinal, the structures appear as elongated linear patterns, the expression was high in all dissected nerves. Furthermore, we quantified $Na_V$1.7 expression in the muscle, heart, spleen, and brain and the signal was negative in these structures of a mouse model. For the kidney, the signal was positive in mice, and for the liver the signal was high, Figure 3a. These results further validate $Na_V$1.7 as a biomarker of peripheral nerves in mice. Figure 3b shows a schematic overview of the approach used to image mouse nerves using fluorescent tracers based on peptide Tsp1a, a venom-derived disulfide-rich peptide with high affinity and selectivity for $Na_V$1.7. For in vivo experiments, the tracers were injected into mice intravenously and were anticipated to delineate nerves. About 30 min after the injection, mice were sacrificed, and nerves were found to retain and accumulate the Tsp1a tracers.

### General Information for Supplementary Information

### ONLINE METHODS

**General.** All solvents, reagents and glassware were obtained from Sigma-Aldrich or Fisher Scientific and used without further purification or cleaning. The azide dyes were commercially accessed or prepared as we previously reported.[1] Acetonitrile (ACN) and water ($H_2O$) were of high-performance liquid chromatography (HPLC) grade and of liquid chromatography mass spectroscopy (LCMS) grade. Phosphate-buffered saline (PBS) without $Ca^{2+}$ or $Mg^{2+}$ was prepared and obtained from the Media Preparation Facility at Memorial Sloan Kettering Cancer Center (MSKCC) and used for all *in vivo* formulations for injections. Reverse-phase (RP) HPLC purifications were performed on a Shimadzu HPLC system (L20154650561) equipped with a DGU-20A degasser, SPD-M20A UV detector, LC-20AB pump system, and a CBM-20A communication BUS module using an Atlantis T3 C18 RP-HPLC column (5 µm, 4.6 x 250 mm, P/N:186003748). Electrospray ionization mass spectroscopy (ESI-MS) spectra were acquired using a Waters Aquity UPLC (Milford, CA) with an electrospray ionization SQ detector. Epifluorescence imaging was performed on an IVIS Spectrum (PerkinElmer) *in vivo* imaging system. Fluorescence stereoscope images were obtained with a Lumar fluorescence stereoscope (SteREO Luma.V12, Zeiss, Jena, Germany). Endoscopic fluorescence images of non-human primates (NHPs) were obtained using an endoscopic fluorescence camera (Quest, manufacturer?). Confocal microscopy images were captured using a Leica SP8 inverted-stand confocal microscope equipped with a tunable white light laser with 470–670 nm range. The microscope was also equipped with a 405 nm diode for detection of Hoechst 33342, an argon laser (with 476 nm, 488 nm, 496 nm and 514 nm laser lines) and a 760 nm laser for near infra-red (NIR) imaging coupled with avalanche photo diode detectors (APDs), which were used for detection of fluorescent Tsp1a conjugates.

**Synthesis of Tsp1a-K4.** In brief, Tsp1a peptide was synthesized using a Liberty Prime microwave peptide synthesizer (CEM corporation, NC, USA) on Rink-amide polystyrene resin to afford an amidated C-terminal.[2] The peptide was simultaneously released from the resin and the side chain protecting groups were removed using trifluoroacetic acid (TFA)/triisopropylsilane (TIPS)/water (48:1:1 v/v/v) for 2.5 h. Trituration of crude Tsp1a was carried out in chilled diethyl ether ($Et_2O$), then the precipitated peptide was separated using solvent A/B (45% v/v ACN, 0.05% v/v TFA), lyophilized, and then pre-purified using C18 RP-HPLC over a period of 1 h. The pure peptide was eluted using a linear gradient of 10–60% solvent B (90% v/v ACN; 0.05% v/v TFA) over 50 min with a flow rate of 8 $mL \cdot min^{-1}$. Oxidation of free cysteines to form the three native disulfide bonds in Tsp1a took place at room temperature for 16 h in a buffer containing 2 M urea,

0.1 M Tris pH 8, 0.15 mM reduced glutathione (GSH), and 0.3 mM oxidized GSH. The purified and oxidized synthetic Tsp1a-K4 was obtained as a white powder. LC-ESI-MS (ES+): m/z calculated for oxidized Tsp1a-K4 – [$C_{148}H_{221}N_{40}O_{40}S_6$] 3389.52; [$C_{148}H_{221}N_{40}O_{40}S_6$+2H]$^{2+}$ 1695.50, observed 1696.00; [$C_{148}H_{221}N_{40}O_{40}S_6$+3H]$^{3+}$ 1130.67, observed 1131.05; [$C_{148}H_{221}N_{40}O_{40}S_6$+4H]$^{4+}$ 848.25, observed 848.50; [$C_{148}H_{220}N_{40}O_{40}S_6$+5H]$^{5+}$ 678.80, observed 679.05.

**Synthesis of Tsp1a-Pra0.** Tsp1a-Pra0 was synthesized using a Liberty Prime microwave peptide synthesizer (CEM corporation, NC, USA) on Rink-amide polystyrene resin to afford an amidated C-terminal. A non-native propargyl glycine was added at the N-terminus (position zero (0)) to provide an attachment point for fluorophores, then the Tsp1a-Pra0 was released from the resin and the side chain protecting groups removed TFA/TIPS/water (48:1:1 v/v/v) for 2.5 h. Trituration of crude Tsp1a-Pra0 was carried out in chilled diethyl ether ($Et_2O$), then the precipitated peptide was separated using solvent A/B (45% v/v acetonitrile, 0.05% v/v TFA), lyophilized, and partially purified using C18 RP-HPLC. The purified Tsp1a-Pra0 was eluted using a linear gradient of 10–80% solvent B (90% v/v ACN; 0.05% v/v TFA) over 50 min with a flow rate of 8 mL·min$^{-1}$. Oxidation of free cysteines took place at room temperature for 16 h in a buffer containing 2 M urea, 0.1 M Tris pH 8, 0.15 mM reduced GSH, and 0.3 mM oxidized GSH. A single peak was obtained from the final analytical RP-HPLC purification and 98% purity was achieved as calculated from the area under the curve. LC-ESI-MS (ES+): m/z calculated for the Tsp1a-Pra0 peptide – [$C_{153}H_{226}N_{42}O_{40}S_6$] 3483.53; [$C_{153}H_{226}N_{42}O_{40}S_6$+2H]$^{2+}$ 1743.00, observed [M+2H]$^{2+}$ 1744.10; [$C_{153}H_{226}N_{42}O_{40}S_6$+3H]$^{3+}$ 1162.33, observed [M+3H]$^{3+}$ 1163.71; [$C_{153}H_{226}N_{42}O_{40}S_6$+4H]$^{4+}$ 872.00, observed [M+4H]$^{4+}$ 872.63; [$C_{153}H_{226}N_{42}O_{40}S_6$+5H]$^{5+}$ 697.80, observed [M+5H]$^{5+}$ 699.00.

**Synthesis of Tsp1a-IR800$_{K4}$.** We conjugated IR800 to lysine 4 (K4) to afford the fluorescently labeled Tsp1a-IR800$_{K4}$ peptide. Tsp1a-K4 peptides (0.37 mM, 250 µg in 100 µL of ACN:water, 50:50) and $Na_2CO_3$ (1 M, 40 µL) were dissolved in 300 µL of a 50:50 mixture of water:ACN inside a 3 mL amber vial with a magnetic bar stirrer. IR800-NHS (0.4 mM, 50 µg in 100 µL of ACN) was dissolved in ACN and added to the reaction mixture dropwise. The reaction mixture was allowed to react for at least 10–30 min. The product of the reaction mixture was separated using HPLC and used for quality control. LC-ESI-MS (ES+): m/z calculated for Tsp1a-IR800$_{K4}$ – [$C_{194}H_{270}N_{43}O_{53}S_{10}$] 4369.70; [$C_{194}H_{270}N_{43}O_{53}S_{10}$+3H]$^{3+}$ 1457.57, observed 1400.67, [$C_{194}H_{270}N_{43}O_{53}S_{10}$+4H]$^{4+}$ 1093.43, observed 1050.75.

**Synthesis of Tsp1a-JA669$_{K4}$.** Tsp1a-K4 (0.37 mM, 250 µg in 100 µL of ACN:water, 50:50) and $Na_2CO_3$ (1 M, 40 µL) were dissolved in 300 µL of a 50:50 mixture of water:ACN inside a 3 mL amber vial with a magnetic bar stirrer. Janelia669-NHS (0.7 mM, 50 µg in 100 µL ACN) was dissolved in ACN and added to the reaction mixture dropwise. The reaction mixture was allowed to react for at least 10–30 min. The products of the reaction mixture were separated using HPLC and used for quality control. LC-ESI-MS (ES+): m/z calculated for Tsp1a-JA669$_{K4}$ – [$C_{178}H_{246}F_3N_{43}O_{42}S_7Si$] 3966.62; [$C_{178}H_{246}F_3N_{43}O_{42}S_7Si+2H]^{2+}$ 1984.31, observed 1985.52; [$C_{178}H_{246}F_3N_{43}O_{42}S_7Si+3H]^{3+}$ 1323.21, observed 1325.02; [$C_{178}H_{246}F_3N_{43}O_{42}S_7Si+4H]^{4+}$ 992.66, observed 993.83.

**Synthesis of Tsp1a-BO665$_{K4}$.** Tsp1a-K4 (0.37 mM, 250 µg in 100 µL of ACN:water, 50:50) and $Na_2CO_3$ (1 M, 40 µL) were dissolved in 300 µL of a 50:50 mixture of water:ACN inside a 3 mL amber vial with a magnetic bar stirrer. BODIPY665-NHS (0.8 mM, 50 µg in 100 µL ACN) was dissolved in ACN and added to the reaction mixture dropwise. The reaction mixture was allowed to react for at least 10–30 min. The products of the reaction mixture were separated using HPLC and used for quality control. LC-ESI-MS (ES+): m/z calculated for Tsp1a-BO665$_{K4}$ – [$C_{177}H_{248}BF_2N_{45}O_{42}S_6$] 3916.70; [$C_{177}H_{248}BF_2N_{45}O_{42}S_6+2H]^{2+}$ 1959.35, observed 1960.50; [$C_{177}H_{248}BF_2N_{45}O_{42}S_6+3H]^{3+}$ 1306.57, observed 1307.89; [$C_{177}H_{248}BF_2N_{45}O_{42}S_6+4H]^{4+}$ 980.18, observed 981.55.

**Synthesis of Tsp1a-DY684$_{K4}$.** Tsp1a-K4 (0.37 mM, 250 µg in 100 µL of ACN:water, 50:50) and $Na_2CO_3$ (1 M, 40 µL) were dissolved in 300 µL of a 50:50 mixture of water:ACN inside a 3 mL amber vial with a magnetic bar stirrer. DY684-NHS (0.4 mM, 50 µg in 100 µL ACN) was dissolved in ACN and added to the reaction mixture dropwise. The reaction mixture was allowed to react for at least 10–30 min. The product of the reaction mixture was separated using HPLC and used for quality control. LC-ESI-MS (ES+): m/z calculated for Tsp1a-DY684$_{K4}$ – [$C_{194}H_{266}N_{43}O_{53}S_{10}$] 4365.67; [$C_{194}H_{266}N_{43}O_{53}S_{10}+3H]^{3+}$ 1456.22, observed 1458.61; [$C_{194}H_{266}N_{43}O_{53}S_{10}+4H]^{4+}$ 1092.42, observed 1093.55.

**Synthesis of Tsp1a-CY7.5$_{K4}$.** Tsp1a-K4 (0.37 mM, 250 µg in 100 µL of ACN:water, 50:50) and $Na_2CO_3$ (1 M, 40 µL) were dissolved in 300 µL of a 50:50 mixture of water:ACN inside a 3 mL amber vial with a magnetic bar stirrer. CY7.5-NHS (0.4 mM, 50 µg in 100 µL ACN) was dissolved in ACN and added to the reaction mixture dropwise. The reaction mixture was allowed to react for at least 10–30 min. The product of the reaction mixture was separated using HPLC and used for quality control. LC-ESI-MS (ES+): m/z calculated for Tsp1a-CY7.5$_{K4}$ – [$C_{193}H_{268}N_{43}O_{40}S_6$] 4019.86; [$C_{193}H_{268}N_{43}O_{40}S_6$+3H]$^{3+}$ 1340.95, observed 1342.09; [$C_{193}H_{268}N_{43}O_{40}S_6$+4H]$^{4+}$ 1005.97, observed 1007.12; [$C_{193}H_{268}N_{43}O_{40}S_6$+5H]$^{5+}$ 804.97, observed 806.15.

**Synthesis of IR800 azide, Janelia669 azide, BODIPY665 azide, Cy7.5 azide and DY684 azide.** IR800, Janelia669, BODIPY665, Cy7.5 and DY684 azides were synthesized by diluting 2 mg (9.2 µmol) 11-Azido-3,6,9-trioxaundecan-1- amine in dichloromethane (0.5 mL) and adding 2 mg of IR800, Janelia669, BODIPY665, Cy7.5 or DY684-NHS esters (1.8 µmol, 2.9 µmol, 3.1 µmol, 1.8 µmol and 2.4 µmol, respectively, with all compounds dissolved in 100 µL ACN just before addition) inside an amber vial. The reaction mixtures were stirred at room temperature in the dark for at least 3 h. The solvents were removed under reduced pressure and the individual crude reaction mixture was subjected to HPLC purification, yielding at least 0.5 mg for each chemical reaction. The final analytical RP-HPLC purifications showed 95–98% purity.

**Synthesis of Tsp1a-IR800$_P$.** Tsp1a-IR800$_P$ was synthesized using Tsp1a-Pra0 (0.71 mM, 250 µg in 100 µL $H_2O$), which was diluted in 100 µL of a 25-mM Tris-buffered aqueous solution. To this reaction mixture, 20 µL of a 50-mM solution of L-ascorbic acid in $H_2O$ and 20 µL of a 50-mM aqueous $CuSO_4$ solution, were added. Immediately, to the same reaction, 104 µg (87 nmol) of IR800 azide in 50 µL of 50:50 ACN/$H_2O$ solution was added, and the reaction mixture was stirred at room temperature in the dark for 4 h. The crude mixture was subjected to HPLC purification, yielding 88 µg Tsp1a-IR800$_P$ (19 nmol; 26%). The final analytical RP-HPLC purification showed 96% purity. LC-ESI-MS (ES+): m/z calculated for Tsp1a-IR800$_P$ – [$C_{207}H_{293}N_{48}O_{57}S_{10}$] 4682.87; [$C_{207}H_{293}N_{48}O_{57}S_{10}$+3H]$^{3+}$ 1563.09, observed 1564.10; [$C_{207}H_{293}N_{48}O_{57}S_{10}$+4H]$^{4+}$ 1172.00, observed 1173.20; [$C_{207}H_{293}N_{48}O_{57}S_{10}$+5H]$^{5+}$ 938.16, observed 939.00.

**Synthesis of Tsp1a-JA669$_P$.** Tsp1a-JA669$_P$ was synthesized using Tsp1a-Pra0 (0.71 mM, 250 µg in 100 µL $H_2O$), which was diluted in 100 µL of a 25-mM Tris-buffered aqueous solution. To this reaction mixture, 20 µL of a 50-mM solution of L-ascorbic acid in $H_2O$ and 20 µL of a 50-mM aqueous $CuSO_4$ solution, were added. Immediately, to the same reaction, 69 µg (87 nmol) of Janelia669 azide in 50 µL of 50:50 ACN/$H_2O$ solution was added, and the reaction mixture was stirred at room temperature in the dark for 4 h. The crude mixture was subjected to HPLC purification, yielding 95 µg of Tsp1a-JA669$_P$ (22 nmol; 31%). The final analytical RP-HPLC purification showed 95% purity. LC-ESI-MS (ES+): m/z calculated for Tsp1a-JA669$_P$ – [$C_{191}H_{269}F_3N_{48}O_{46}S_7Si$] 4279.80; [$C_{191}H_{269}F_3N_{48}O_{46}S_7Si+3H]^{3+}$ 1427.60, observed 1428.80; [$C_{191}H_{269}F_3N_{48}O_{46}S_7Si+4H]^{4+}$ 1070.95, observed 1072.20; [$C_{191}H_{269}F_3N_{48}O_{46}S_7Si+5H]^{5+}$ 856.96, observed 857.97.

**Synthesis of Tsp1a-BO665$_P$.** Tsp1a-BO665$_P$ was synthesized using Tsp1a-Pra0 (0.71 mM, 250 µg in 100 µL $H_2O$), which was diluted in 100 µL of a 25-mM Tris-buffered aqueous solution. To this reaction mixture, 20 µL of a 50-mM solution of L-ascorbic acid in $H_2O$ and 20 µL of a 50-mM aqueous $CuSO_4$ solution, were added. Immediately, to the same reaction, 65 µg (87 nmol) of BODIPY665 azide in 50 µL of 50:50 ACN/$H_2O$ solution was added, and the reaction mixture was stirred at room temperature in the dark for 4 h. The crude mixture was subjected to HPLC purification, yielding 70 µg of Tsp1a-BO665$_P$ (17 nmol; 23%). The final analytical RP-HPLC purification showed 96% purity. LC-ESI-MS (ES+): m/z calculated for Tsp1a-BO665$_P$ – [$C_{190}H_{271}BF_2N_{50}O_{46}S_6$] 4229.88; [$C_{190}H_{271}BF_2N_{50}O_{46}S_6+3H]^{3+}$ 1410.96, observed 1412.22; [$C_{190}H_{271}BF_2N_{50}O_{46}S_6+4H]^{4+}$ 1058.47, observed 1059.66; [$C_{190}H_{271}BF_2N_{50}O_{46}S_6+5H]^{5+}$ 846.98, observed 848.25.

**Synthesis of Tsp1a-DY684$_P$.** Tsp1a-DY684$_P$ was synthesized using Tsp1a-Pra (0.71 mM, 250 µg in 100 µL $H_2O$), which was diluted in 100 µL of a 25-mM Tris-buffered aqueous solution. To this reaction mixture, 20 µL of a 50-mM solution of L-ascorbic acid in $H_2O$ and 20 µL of a 50-mM aqueous $CuSO_4$ solution, were added. Immediately, to the same reaction, 103 µg (86 nmol) of DY684 azide in 50 µL of 50:50 ACN/$H_2O$ solution was added, and the reaction mixture was stirred at room temperature in the dark for 4 h. The crude mixture was subjected to HPLC purification, yielding 75 µg of Tsp1a-DY684$_P$ (16 nmol; 22%). The final analytical RP-HPLC purification showed 95% purity. LC-ESI-MS (ES+): m/z calculated for Tsp1a-DY684$_P$ – [$C_{207}H_{289}N_{48}O_{57}S_{10}$] 4678.84; [$C_{207}H_{289}N_{48}O_{57}S_{10}+3H]^{3+}$ 1560.61, observed 1562.72; [$C_{207}H_{289}N_{48}O_{57}S_{10}+4H]^{4+}$ 1170.71, observed 1171.82; [$C_{207}H_{289}N_{48}O_{57}S_{10}+5H]^{5+}$ 936.77, observed 937.08.

**Synthesis of Tsp1a-Cy7.5$_P$.** Tsp1a-Cy7.5$_P$ was synthesized using Tsp1a-Pra0 (0.71 mM, 250 µg in 100 µL $H_2O$), which was diluted in 100 µL of a 25-mM Tris-buffered aqueous solution. To this reaction mixture, 20 µL of a 50-mM solution of L-ascorbic acid in $H_2O$ and 20 µL of a 50-mM aqueous $CuSO_4$ solution, were added. Immediately, to the same reaction, 73 µg (86 nmol) of

Cy7.5 azide in 50 µL of 50:50 (ACN/$H_2O$) solution was added, and the reaction mixture was stirred at room temperature in the dark for 4 h. The crude mixture was subjected to HPLC purification, yielding 90 µg of Tsp1a-Cy7.$5_P$ (21 nmol; 29%). The final analytical RP-HPLC purification showed 96% purity. LC-ESI-MS (ES+): m/z calculated for Tsp1a-Cy7.$5_P$ – [$C_{206}H_{291}N_{48}O_{44}S_6$] 4333.03; [$C_{206}H_{291}N_{48}O_{44}S_6$+3H]$^{3+}$ 1445.34, observed 1446.30; [$C_{206}H_{291}N_{48}O_{44}S_6$+4H]$^{4+}$ 1084.26, observed 1085.44; [$C_{206}H_{291}N_{48}O_{44}S_6$+5H]$^{5+}$ 867.61, observed 868.87.

**Tryptic digest of Tsp1a-K4 peptides and Tsp1a**

15 µL of digestion buffer and 1.5 µL of reducing buffer were added to a 0.5 mL microcentrifuge tube (MT) at room temperature. 5 µg of Tsp1a-K4 tracers were dissolved in 10.5 µL of a solution which was added to the same MT, then the final volume was adjusted to 28 µL with ultrapure water, if needed. The sample was then incubated at 95 °C for 5 min. The sample was allowed to cool to room temperature, then 2 µL of activated trypsin was immediately added to the MT, followed by incubation at 37 °C overnight. LC-ESI-MS ($ES^+$) after the digestions: For Tsp1a-IR$800_{K4}$ m/z – [$C_{158}H_{216}N_{33}O_{47}S_9$] 3444.33; [$C_{152}H_{213}N_{33}O_{43}S_8$+2H]$^{2+}$ 1723.17, observed 1723.32; [$C_{152}H_{213}N_{33}O_{43}S_8$+3H]$^{3+}$ 1149.17, observed 1149.41.

For Tsp1a-JA$669_{K4}$ – [$C_{140}H_{188}F_3N_{33}O_{36}S_6$] 3156.22; [$C_{140}H_{188}F_3N_{33}O_{36}S_6$+2H]$^{2+}$ 1579.11, observed 1580.02, [$C_{140}H_{188}F_3N_{33}O_{36}S_6$+3H]$^{3+}$ 1053.07, observed 1053.81.

For Tsp1a-BO$665_{K4}$ – [$C_{141}H_{194}BF_2N_{35}O_{36}S_5$] 3113.3; [$C_{141}H_{194}N_{35}O_{36}S_5$+2H]$^{2+}$ 1557.65, observed 1558.12; [$C_{141}H_{194}N_{35}O_{36}S_5$+3H]$^{3+}$ 1038.77, observed 1038.91.

For Tsp1a-DY$684_{K4}$ – [$C_{158}H_{212}F_3N_{33}O_{47}S_9$] 3611.27; [$C_{158}H_{212}F_3N_{33}O_{47}S_9$+2H]$^{2+}$ 1806.64, observed 1807.24; [$C_{158}H_{212}F_3N_{33}O_{47}S_9$+3H]$^{3+}$ 1204.76, observed 1204.59.

For Tsp1a-CY7.$5_{K4}$ – [$C_{157}H_{214}N_{33}O_{34}S_5$] 3265.46; [$C_{157}H_{214}N_{33}O_{34}S_5$+2H]$^{2+}$ 1633.73, observed 1633.95; [$C_{157}H_{214}N_{33}O_{34}S_5$+3H]$^{3+}$ 1089.49, observed 1089.92.

LC-ESI-MS analysis of the tryptic digestion of Tsp1a revealed the following fragments: [$C_{89}H_{135}N_{25}O_{27}S_4$] 1058.45, observed 1058.66; [$C_{89}H_{135}N_{25}O_{27}S_4$] 705.97, observed 706.33.

**Cell Lines.** HEK293 cells stably expressing the human $Na_V$ channel β1 subunit (hNaVβ1) in combination with the α subunit of $hNa_V1.7$ (Scottish Biomedical, Glasgow, UK) were seeded into a 175 $cm^2$ cell culture flask two days prior to patching and were detached at 60% confluency using

2 mL Detachin™ (Genlantin, San Diego, CA, USA). After pelleting, cells were centrifuged at 800 rpm for 8 min, then the supernatant was removed and cells were resuspended in 5 mL QPatch media containing 96.5% CD293 medium, 25 mM HEPES (Gibco), and glutamine (Gibco).

**Electrophysiology.** Whole-cell patch-clamp experiments were performed at room temperature using a QPatch 16X automated electrophysiology platform (Sophion Bioscience, Denmark) using 16-channel planar patch-chip plates (QPlates) with a patch-hole diameter of 1 µm and resistance of 2 MΩ. Whole-cell currents were filtered at 5 kHz (8-pole Bessel) and digitized at 25 kHz. A P4 online leak-subtraction protocol was used with non-leak-subtracted currents acquired in parallel. The extracellular solution was 2 mM $CaCl_2$, 1 mM $MgCl_2$, 10 mM HEPES, 4 mM KCl, 145 mM NaCl, pH adjusted to 7.4 with NaOH, and 294 mOsm. The intracellular solution was 140 mM CsF, 1 mM/5 mM EGTA/CsOH, 10 mM HEPES, 10 mM NaCl, 10 sucrose, pH adjusted to 7.4 with CsOH, and 304 mOsm. Tsp1a and fluorescently-labeled derivatives were dissolved in extracellular solution with 0.1% bovine serum albumin (BSA). Concentration-response data were obtained using eight concentrations of peptide (1, 3, 10, 30, 100, 300, 1000 and 3000 nM). HEK293-h$Na_V$ cells were clamped at a holding potential of –80 mV, then for each concentration, 10 µL of peptide was added for 6 s before applying the following voltage protocol: –80 mV for 10 ms, –120 mV for 200 ms, 0 mV for 20 ms, then return to the holding potential of –80 mV. The voltage protocol was applied 15 times, with cells clamped at a holding potential of –80 mV for 30 s between runs. Concentration-response data were analysed in Prism 8 (GraphPad Software) to obtain $IC_{50}$ values (i.e., the concentration for 50% inhibition of channel currents).

**Human tissue.** All dissection experiments were performed in accordance with institutional guidelines and approved by the MSKCC Review Board. Biomaterial from six human cadavers (men = 3, women = 3) was acquired from Worldwide Primates (Miami, FL, USA), and dissected at the MSKCC necropsy room by trained physicians and necropsy technicians. Nerves (287), muscle, artery and tendons were dissected from the extremities. Nerves (245) and internal organs were dissected from the torsos. All specimens obtained at necropsy were fixed in 4% paraformaldehyde (PFA), and paraffin embedded. Sections of 10-µm thickness from each tissue were submitted for hematoxylin and eosin (H&E) staining and $Na_V$1.7 immunohistochemistry (IHC).

**Animal models.** All animal experiments were performed in accordance with institutional guidelines and approved by the IACUC of MSKCC, following NIH guidelines for animal welfare. All mice used in this study (n = 120 males and 18 females) were athymic nude, between 4 and

10 weeks old, purchased from Envigo (Stock#:069); RMS, INC., USA). All mice were allowed to acclimatize at the MSKCC vivarium for 1 week with food and water available *ad libitum* prior to any experimental procedure. The non-human primates (grivets) used in the study (*Chlorocebus aethiops*, n = 2 males and 2 females) were acquired from Worldwide Primates (Miami, FL, USA) and were between 4 and 6 years old and weighted 4.5–6.5 kg, Grivets were quarantined for 10 weeks and allowed to acclimatize at the MSKCC vivarium for 6 months before the experimental procedure.

**Immunohistochemistry.** Mice, non-human primate (NHP, and human tissues were submitted for IHC at the Molecular Cytology Core Facility of MSKCC. Briefly, paraffin-embedded formalin-fixed 10-μm sections were deparaffinized with EZPrep buffer. An automated method was performed using the Discovery XT processor (Ventana Medical System, Tucson, AZ). Primary anti-$Na_V1.7$ antibody [N68/6] (Abcam ab85015) specifically bound to mouse, NHP, and human $Na_V1.7$ (all using a 0.5 μg/mL concentration). The primary antibody was incubated for 1 h, followed by 1 h incubation with biotinylated goat anti-rabbit IgG (0.8 μg/mL) at a 10x dilution. For expression detection, a 3,3'-diaminobenzidine (DAB) detection kit (Ventana Medical Systems) was used according to the manufacturer's instructions. The sections were then counterstained with hematoxylin and cover slipped with Permount (Fisher Scientific, Pittsburgh, PA). Incubating with a rabbit IgG instead of the primary antibody controlled for non-specific binding of the secondary antibody. Slides were further scanned (Mirax, 3DHISTECH, Budapest, Hungary) to allow for digital histological correlation. Quantification of $Na_V1.7$ expression was carried out on digitalized slides, using the axial sections of the nerve bundles. CaseViewer 2.4.0.119028 software was used to delineate regions of interest over the nerve bundles. Thresholding was performed using a previously described Fiji (Image J) plugin (50) on brown tissue areas ($Na_V1.7$ marked with DAB) and the remaining blue tissue areas. The relative $Na_V1.7$-positive area was calculated by dividing the brown area (DAB) by the blue area (total tissue).

**Confocal microscopy.** In general, staining with fluorescent Tsp1a compounds was performed in both mouse peripheral nerves and human vagus nerves. For mice, 10-μm cryosections of OCT-embedded sciatic nerve tissues from mice previously injected with fluorescent Tsp1a compounds (1 nmol; 10 μM of fluorescent Tsp1a in 100 μL PBS), Tsp1a/fluorescent Tsp1a (fluorescent Tsp1a, 10 μM, 1 nmol and 'cold' Tsp1a, 204 μM, 21 nmol in 100 μL PBS) or PBS, were used for the microscopy experiments. Tissue nuclei were stained with Hoechst 33342 (blue, 20 μM, 1 nmol in 50 μL PBS) up to 90 min post-mortem and placed directly on a microscope slide for imaging. For human tissue, 10-μm cryosections of vagus nerve were immersed in fluorescent Tsp1a (3 nmol,

30 µM of fluorescent Tsp1a in 100 µL PBS), Tsp1a/fluorescent Tsp1a (fluorescent Tsp1a, 30 µM, 3 nmol and 'cold' Tsp1a, 90 µM, 9 nmol in 100 µL PBS) or PBS. Tissues were incubated with Hoechst 33342 (blue, 20 µM, 1 nmol in 50 µL of PBS). For human vagus nerve tissue, the specimen was incubated for at least 2 min, followed by one PBS wash cycle. For the block experiment, nerve tissue was first immersed in Tsp1a solution for at least 5 min, followed by immersion in fluorescent Tsp1a for at least 2 min, followed by one PBS wash cycle.

**Epifluorescence imaging.** In general, animals were intravenously injected with fluorescent Tsp1a (1 nmol, 10 µM of fluorescent Tsp1a in 100 µL PBS, n = 6). To assess the specificity of the fluorescent Tsp1a accumulation, we injected a combination of fluorescent Tsp1a (fluorescent Tsp1a, 10 µM, 1 nmol) and an excess of 'cold' Tsp1a (204 µM, 21 nmol in 100 µL PBS) (n = 6) or PBS (n = 9). Epifluorescence images were obtained in vivo after 5–8 min on the upper limb, lower legs, and spinal cord. Animals were sacrificed 30 min post-injection and epifluorescence images obtained. Epifluorescence images of right sciatic nerves (RSN) and left sciatic nerves (LSN) were obtained first *in situ* and then *ex vivo*, together with excised muscle, heart, kidney, liver and brain tissue using an IVIS Spectrum in vivo imaging system (PerkinElmer) with predefined filter set ranges (excitation = 600/20–780/20 nm, emission = 650–820 nm). Autofluorescence was removed via spectral unmixing. Semiquantitative analysis of the epifluorescence images was conducted by measuring the average radiant efficiency (in units of [p/s/cm$^2$/sr]/[µW/cm$^2$]) in regions of interest (ROIs) that were drawn on all resected organs under white light guidance.

**Fluorescence Stereoscope Imaging.** For fluorescence imaging experiments, animals were sacrificed 30 min after tail vein injection of either fluorescent Tsp1a tracer, block formulation or PBS. The fluorescentTsp1a tracer signal in mouse sciatic nerves was also visualized using a fluorescence stereoscope 30 min after of intravenous injection of fluorescent Tsp1a (1 nmol, 10 µM of Tsp1a in 100 µL PBS, n = 6) or PBS (n = 3/group). Fluorescence images were obtained using mice with exposed but otherwise intact sciatic nerves. Images were also obtained from excised sciatic nerves and muscle. Imaging was performed in bright field and fluorescence mode, with a 600/20–780/20 nm laser excitation and 650–820 nm emission filter and an exposure time of 200–400 ms.

**Estimation of Tissue Penetrance.** A 10 cm phantom mold was prepared and 1 cm-sliced for tissue-mimicking penetration experiments. A clear cylindric hollow membrane tubing (a 20 mL syringe, diameter = 2 cm, length = 14 cm) was chosen to be the main mold-type container to grow the soft-tissue-mimicking material for the phantoms. The soft-tissue-mimicking phantom was freshly prepared by adding 15% v/v intralipid 20% I.V. (Fresenius Kabi AB) fat emulsion (to add scattering feature) and 0.01 mM direct red 81 (for absorption) to a prewarmed solution of 5% v/v agarose type 1 (solid < 37 °C) in Milli-Q water (Millipore Corp., 18.2 MΩ $cm^{-1}$ at 25 °C). At 15% agarose, the phantom was suspended in water, with the weight of the hollow tubing being supported. The mixture was poured into the mold and allowed to cool and solidify for at least 2 h. The phantom was then removed carefully from the syringe and sliced transversally at 1 cm. The phantoms were used for tissue penetration experiments with fluorescent Tsp1a peptides.

**Residence Time (with clinical stereoscope).** The Tsp1a-$IR800_P$ fluorescence signal in mouse peripheral nerves was visualized using a fluorescence stereoscope after intravenous injection with Tsp1a-$IR800_P$ (1 nmol, 9.7 µM of Tsp1a-$IR800_P$ in 100 µL PBS) or PBS (n = 3/group). Fluorescence images were obtained using mice with exposed but otherwise intact peripheral nerves. Imaging was performed in bright field and fluorescence mode, with an 800/20 nm laser excitation and 820/30 emission filter and an exposure time 30–180 min.

**Tsp1a-$IR800_P$ serum half-life and imaging after intravenous injection in NHPs.** Anesthesia was induced in grivets (2 males and 2 females, 4.5–6.5 kg) using isoflurane inhalation under veterinarian supervision (1–5% in $O_2$). The NHP was placed on a table in the supine position with neck extension using a cushion under the scapula. The animal was monitored during the entire procedure and vital signs (blood pressure, heart rate, oxygen saturation, and body temperature) were acquired during the entire anesthesia period, from time zero up to 120 min (experimental endpoint). An intravenous catheter was placed in right the cephalic vein, where Tsp1a-$IR800_P$ (250 µg $kg^{-1}$, in 5 mL PBS) was administered intravenously (5 mL in ~60 s), followed by a 5 mL saline flush. Another catheter was placed in the left cephalic vein for blood draws. A urine catheter was placed in the urethra for urine collection. Blood and urine samples (both 1.0 mL) were taken at predetermined time points (0, 3, 5, 10, 15, 30, 60, 90 and 120 min). A left thyroidectomy was performed to expose the recurrent laryngeal and vagus nerves. A fluorescent camera (Quest Medical Imaging) was placed at 18 cm distance from the neck and videos were acquired before and after contrast application using the Quest Spectrum platform (Quest Medical Imaging) up to 120 min post-injection. The same instrument settings were used throughout the imaging procedure (100% laser power, 25.5 dB gain, and white light exposure 30 ms), except for the laser exposure time which varied between animals (animal 1: 10 ms, animal 2: 40 ms, animal 3: 30 ms, animal 4: 30ms). The NHP was then euthanized with an intravenous overdose of pentobarbital sodium and phenytoin sodium (440 mg $mL^{-1}$). The recurrent laryngeal and vagus nerves were resected for further analysis and stored in ice-cold PBS until imaging. The freshly excised nerves along with the thyroid (right lobe pre-injection and left lobe post-injection), and a fragment of the pre-thyroid muscles (also pre- and post-injection) were imaged using an IVIS Spectrum (Zeiss, Germany) to identify Tsp1a-IR800$_P$ signal in the nerves. We also imaged 100 µL of blood and 100 µL of urine at the different time points.

For blood and urine metabolite analysis, a 1:1 mixture of ACN:dimethyl sulfoxide (1.0 mL) was added to blood samples (1.0 mL) inside Eppendorf tubes and then vortexed for 2 min at room temperature. Samples were centrifuged at 4500 rpm (4° C) for 6 min, then the supernatant was removed and placed into new Eppendorf tubes. These Eppendorf tubes were left to stand for 30 min, then the supernatant was removed and analyzed on LCMS with 95% ACN/5% $H_2O$/0.1% formic acid.

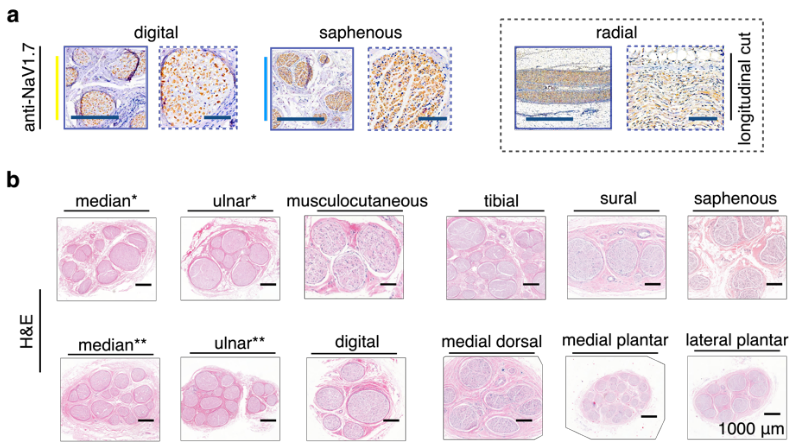


**Figure S1.** Expression of $Na_V1.7$ in human peripheral nerves. (a) Representative IHC staining performed on human peripheral nervous system to demonstrate the high expression of sodium $Na_V1.7$ in human nerves. These human peripheral nerve samples were also sliced longitudinally before staining. (b) Adjacent H&E slices of resected human peripheral nerves showing nerve bundles and circular patterns for nerves cut transversally. These representative human nerves included median (* = arm, ** = hand), ulnar (* = arm, ** = hand), musculocutaneous, tibial, sural, saphenous, digital, medial dorsal, medial plantar and lateral plantar.

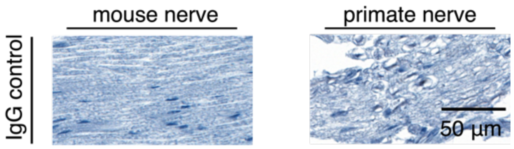


**Figure S2.** Immunoglobulin (IgG) antibody control for resected peripheral nerve samples. Immunohistochemical IgG control was used in resected mouse and NHP peripheral nerve samples alongside anti-$Na_V$1.7 IHC to validate $Na_V$1.7 expression.

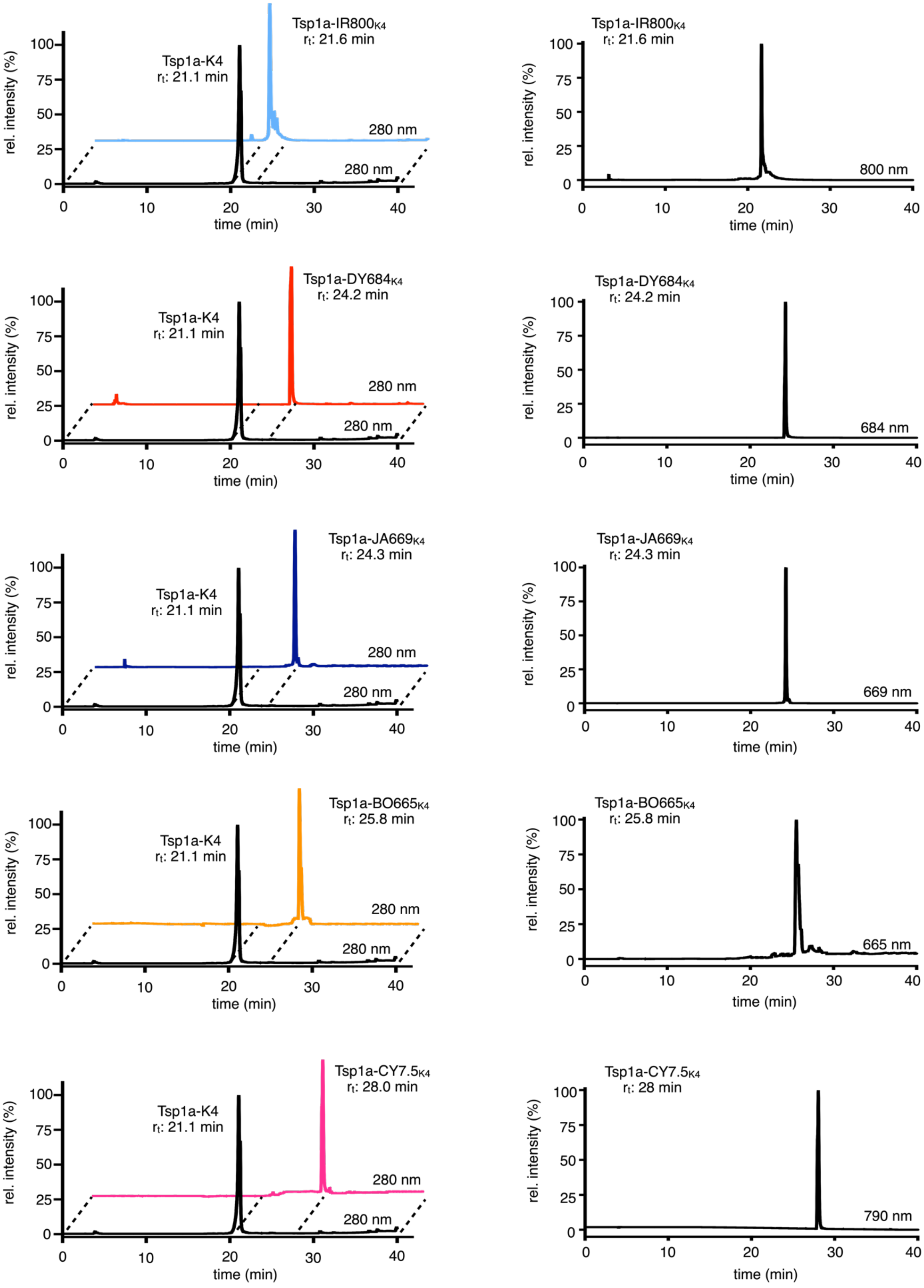

Tsp1a-K4
$r_t$: 21.1 min
Tsp1a-IR800$_{K4}$
$r_t$: 21.6 min
280 nm
280 nm
800 nm
Tsp1a-DY684$_{K4}$
$r_t$: 24.2 min
684 nm
Tsp1a-JA669$_{K4}$
$r_t$: 24.3 min
669 nm
Tsp1a-BO665$_{K4}$
$r_t$: 25.8 min
665 nm
Tsp1a-CY7.5$_{K4}$
$r_t$: 28.0 min
$r_t$: 28 min
790 nm
rel. intensity (%)
time (min)

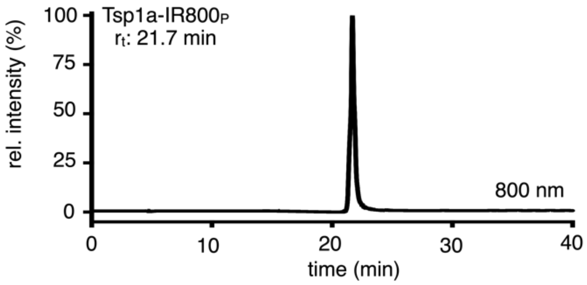
Tsp1a-IR800P
rt: 21.7 min
800 nm
rel. intensity (%)
time (min)

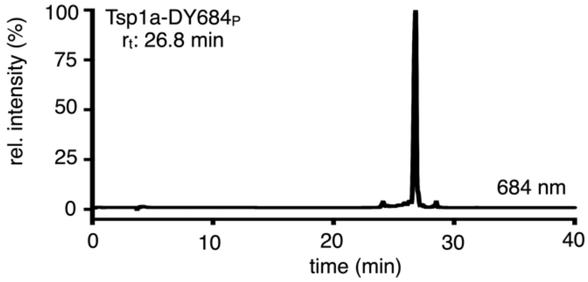
Tsp1a-DY684P
rt: 26.8 min
684 nm
rel. intensity (%)
time (min)

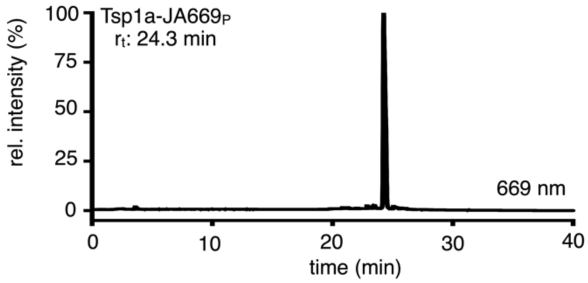
Tsp1a-JA669P
rt: 24.3 min
669 nm
rel. intensity (%)
time (min)

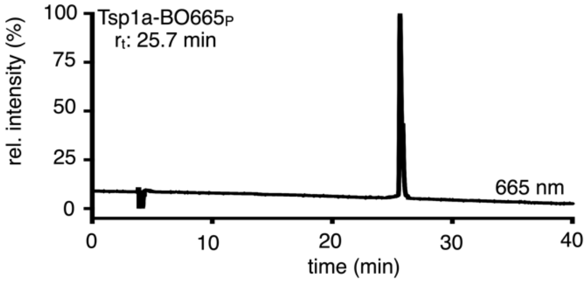
Tsp1a-BO665P
rt: 25.7 min
665 nm
rel. intensity (%)
time (min)

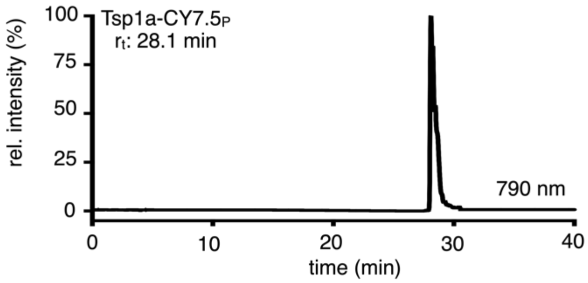
Tsp1a-CY7.5P
rt: 28.1 min
790 nm
rel. intensity (%)
time (min)

**Figure S3.** Chemical features of fluorescently labeled Tsp1a peptides. RP-HPLC chromatograms showing elution of Tsp1a-K4, Tsp1a-IR800$_{K4}$, Tsp1a-JA669$_{K4}$, Tsp1a-BO665$_{K4}$, Tsp1a-DY684$_{K4}$ and Tsp1a-CY7.5$_{K4}$ peptides (in black, light blue, red, blue, orange and pink, respectively), with peptide absorbance measured at 280 nm (left) and chromatograms at right showing the characteristic absorbance of the covalently attached fluorophores IR800 (800 nm), DY684 (670 nm), Janelia669 (670 nm), Bodipy665 (670 nm) and CY7.5 (790 nm).

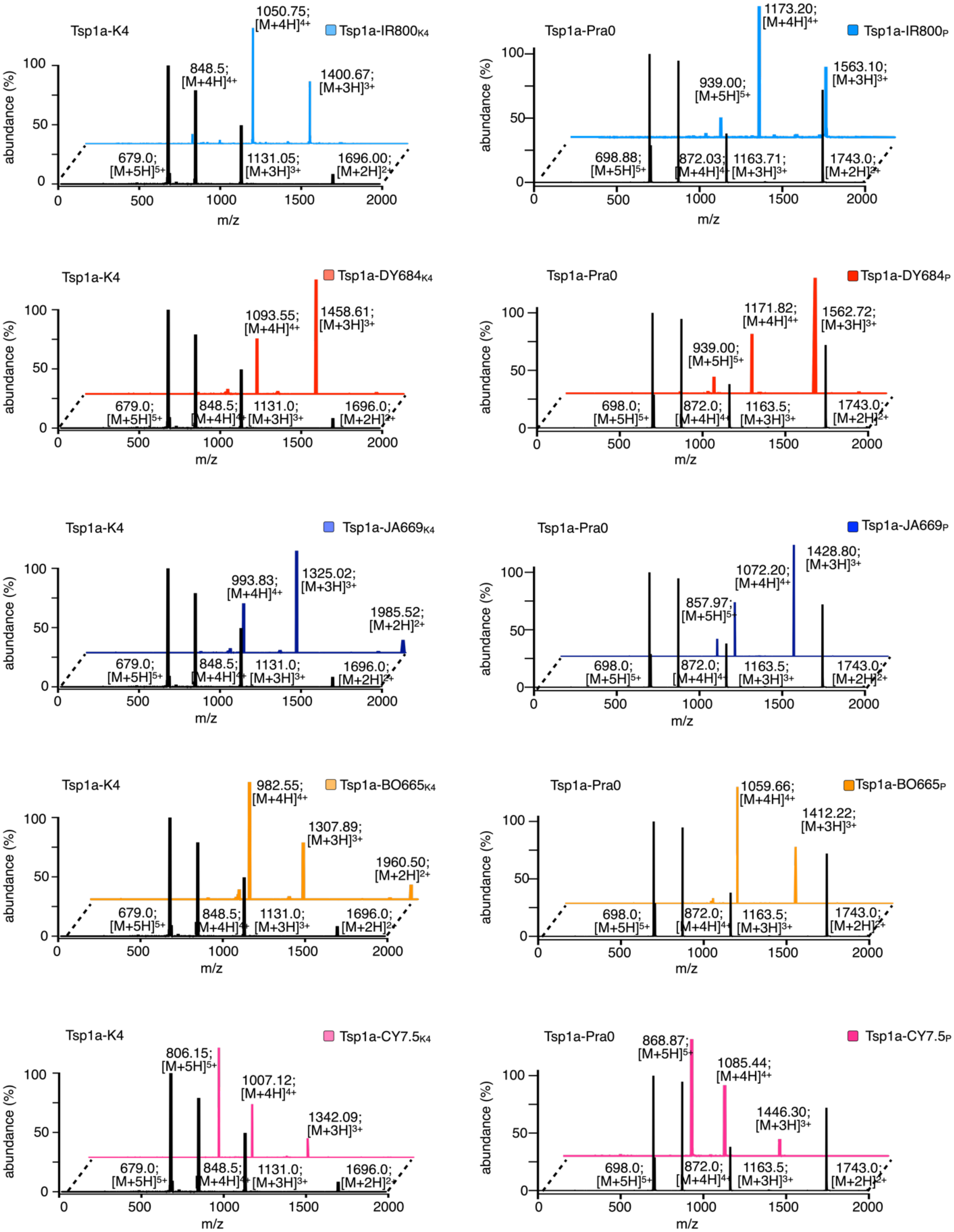

Tsp1a-K4
Tsp1a-IR800$_{K4}$
1050.75; [M+4H]$^{4+}$
848.5; [M+4H]$^{4+}$
1400.67; [M+3H]$^{3+}$
679.0; [M+5H]$^{5+}$
1131.05; [M+3H]$^{3+}$
1696.00; [M+2H]$^{2+}$
Tsp1a-Pra0
Tsp1a-IR800$_{P}$
1173.20; [M+4H]$^{4+}$
1563.10; [M+3H]$^{3+}$
939.00; [M+5H]$^{5+}$
698.88; [M+5H]$^{5+}$
872.03; [M+4H]$^{4+}$
1163.71; [M+3H]$^{3+}$
1743.0; [M+2H]$^{2+}$
Tsp1a-K4
Tsp1a-DY684$_{K4}$
1093.55; [M+4H]$^{4+}$
1458.61; [M+3H]$^{3+}$
679.0; [M+5H]$^{5+}$
848.5; [M+4H]$^{4+}$
1131.0; [M+3H]$^{3+}$
1696.0; [M+2H]$^{2+}$
Tsp1a-Pra0
Tsp1a-DY684$_{P}$
1171.82; [M+4H]$^{4+}$
1562.72; [M+3H]$^{3+}$
939.00; [M+5H]$^{5+}$
698.0; [M+5H]$^{5+}$
872.0; [M+4H]$^{4+}$
1163.5; [M+3H]$^{3+}$
1743.0; [M+2H]$^{2+}$
Tsp1a-K4
Tsp1a-JA669$_{K4}$
993.83; [M+4H]$^{4+}$
1325.02; [M+3H]$^{3+}$
1985.52; [M+2H]$^{2+}$
679.0; [M+5H]$^{5+}$
848.5; [M+4H]$^{4+}$
1131.0; [M+3H]$^{3+}$
1696.0; [M+2H]$^{2+}$
Tsp1a-Pra0
Tsp1a-JA669$_{P}$
1428.80; [M+3H]$^{3+}$
1072.20; [M+4H]$^{4+}$
857.97; [M+5H]$^{5+}$
698.0; [M+5H]$^{5+}$
872.0; [M+4H]$^{4+}$
1163.5; [M+3H]$^{3+}$
1743.0; [M+2H]$^{2+}$
Tsp1a-K4
Tsp1a-BO665$_{K4}$
982.55; [M+4H]$^{4+}$
1307.89; [M+3H]$^{3+}$
1960.50; [M+2H]$^{2+}$
679.0; [M+5H]$^{5+}$
848.5; [M+4H]$^{4+}$
1131.0; [M+3H]$^{3+}$
1696.0; [M+2H]$^{2+}$
Tsp1a-Pra0
Tsp1a-BO665$_{P}$
1059.66; [M+4H]$^{4+}$
1412.22; [M+3H]$^{3+}$
698.0; [M+5H]$^{5+}$
872.0; [M+4H]$^{4+}$
1163.5; [M+3H]$^{3+}$
1743.0; [M+2H]$^{2+}$
Tsp1a-K4
Tsp1a-CY7.5$_{K4}$
806.15; [M+5H]$^{5+}$
1007.12; [M+4H]$^{4+}$
1342.09; [M+3H]$^{3+}$
679.0; [M+5H]$^{5+}$
848.5; [M+4H]$^{4+}$
1131.0; [M+3H]$^{3+}$
1696.0; [M+2H]$^{2+}$
Tsp1a-Pra0
Tsp1a-CY7.5$_{P}$
868.87; [M+5H]$^{5+}$
1085.44; [M+4H]$^{4+}$
1446.30; [M+3H]$^{3+}$
698.0; [M+5H]$^{5+}$
872.0; [M+4H]$^{4+}$
1163.5; [M+3H]$^{3+}$
1743.0; [M+2H]$^{2+}$
abundance (%)
m/z

**Figure S4.** LC-MS spectra of Tsp1a-K4, Tsp1a-IR800$_{K4}$, Tsp1a-JA669$_{K4}$, Tsp1a-BO665$_{K4}$, Tsp1a-DY684$_{K4}$ and Tsp1a-CY7.5$_{K4}$ peptides (left) and Tsp1a-Pra0, Tsp1a-IR800$_{P}$, Tsp1a-JA669$_{P}$, Tsp1a-BO665$_{P}$, Tsp1a-DY684$_{P}$ and Tsp1a-CY7.5$_{P}$ peptides (right). These tracers show major ion species that correspond to the calculated mass of the synthetic fluorescently labeled Tsp1a-K4 and Tsp1a-Pra0 peptides.

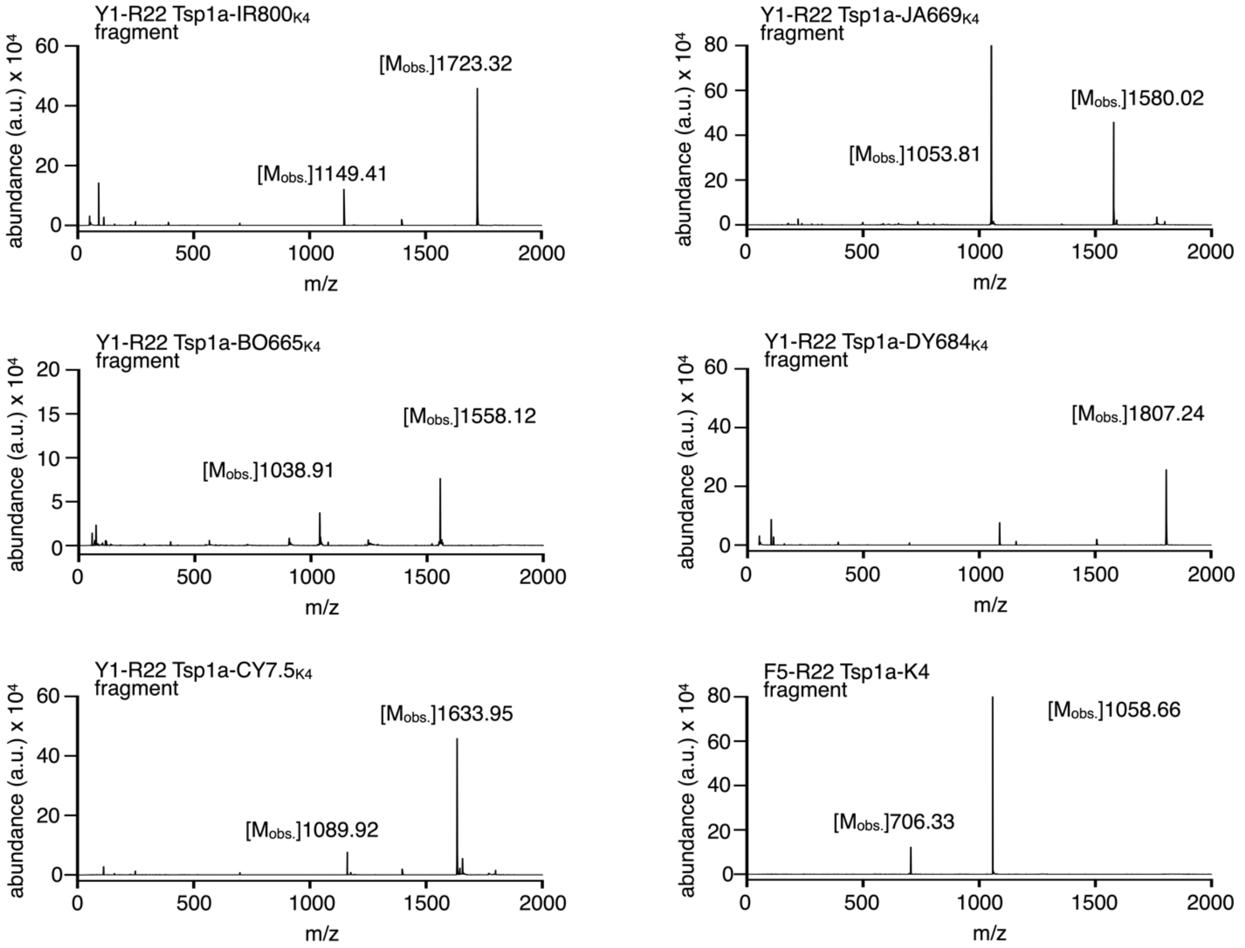


**Figure S5.** Mas spectrometric analysis of tryptic digestion(s) of Tsp1a-K4 tracers and Tsp1a-K4. In general, two major tryptic fragments were observed in all the digestions. The Y1-R22 fragment of fluorescent Tsp1a-K4 was found coupled to the corresponding fluorophore. Tryptic digestion of Tsp1a-K4 led to a F5-R22 fragment.

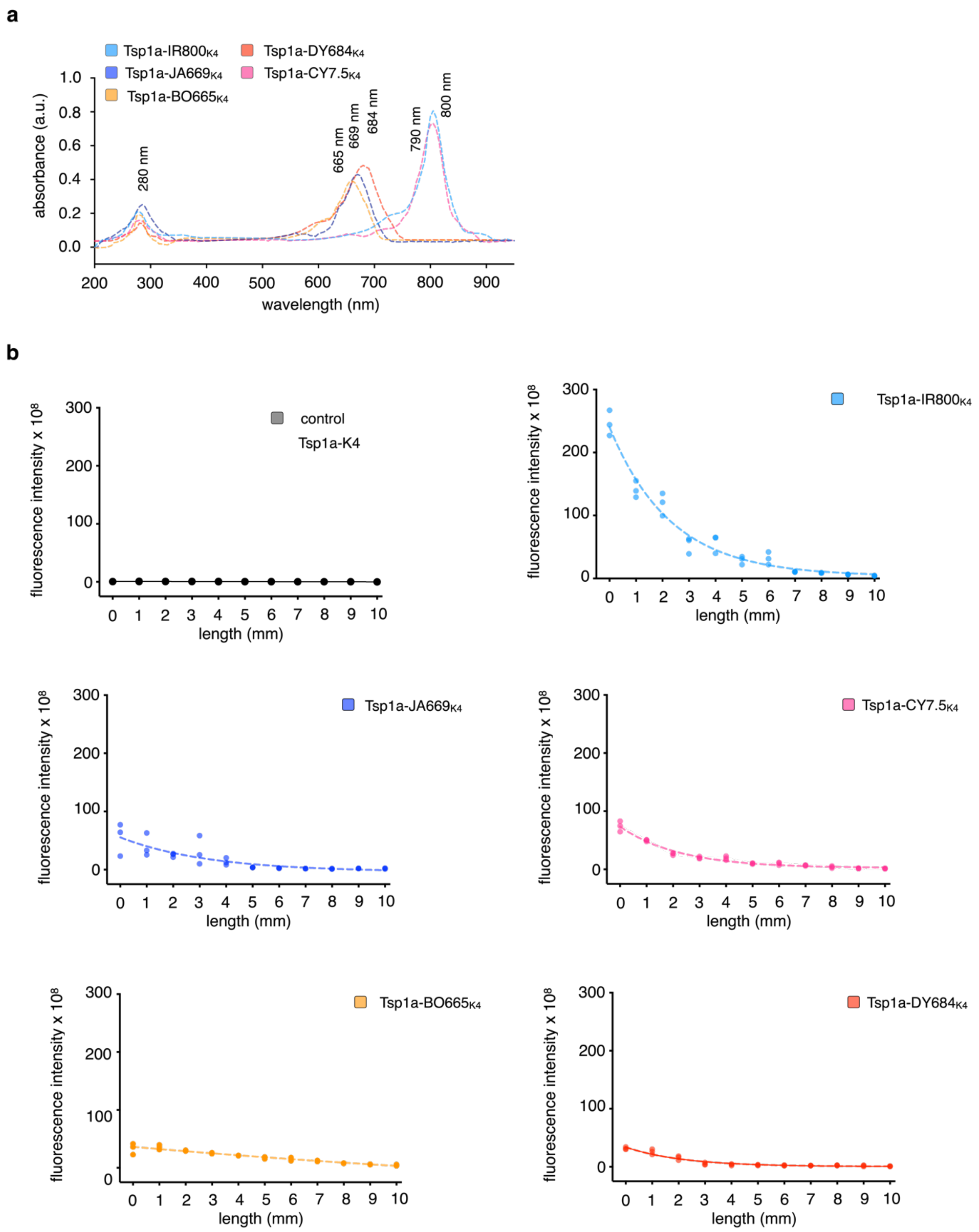
a
Tsp1a-IR800K4
Tsp1a-JA669K4
Tsp1a-BO665K4
Tsp1a-DY684K4
Tsp1a-CY7.5K4
absorbance (a.u.)
wavelength (nm)
280 nm
665 nm
669 nm
684 nm
790 nm
800 nm
b
control
Tsp1a-K4
fluorescence intensity x 10^8
length (mm)

**Figure S6.** Photochemical features of the Tsp1a peptides. (a) Absorbance spectra of 0.2 μM Tsp1a-Pra0, Tsp1a-IR800$_P$, Tsp1a-JA669$_P$, Tsp1a-BO665$_P$, Tsp1a-DY684$_P$ and Tsp1a-CY7.5$_P$ pptides (in black, light blue, blue, orange, red and pink, respectively) observed from 200–900 nm. Classical absorbance maxima were observed for each of the fluorophores. (b) Fluorescence intensity of fluorescently labeled Tsp1a-K4 peptides in the presence of synthetic phantoms which helped to determine the penetrance of each Tsp1a tracer.

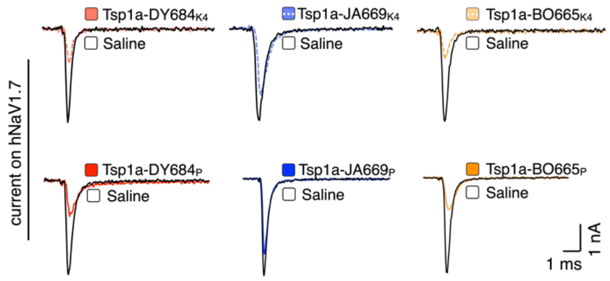


**Figure S7.** Representative $hNa_V1.7$ currents in the absence (black trace) or presence (colored traces) of 5 μM Tsp1a tracer confirming the ability of each peptide to inhibit $hNa_V1.7$.

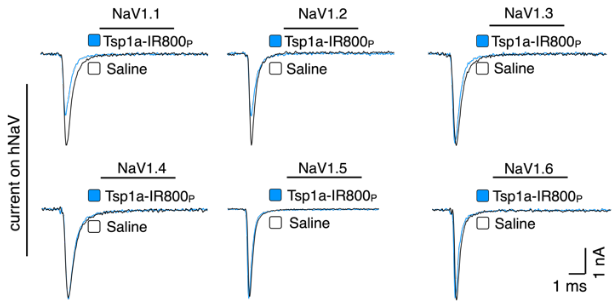


**Figure S8.** Representative currents from a panel of human $Na_V$ channels in the absence (black traces) or presence (blue traces) of 5 μM Tsp1a-$IR800_{P.}$ Tsp1a-$IR800_P$ has no effect on $hNa_V1.2$–1.6 and it causes only minor inhibition of $Na_V1.1$.

| * peptide acronym | Dye (Z) | Absorbance max.$^{DMSO}$ | Emission max.$^{DMSO}$ | Yield [%] | Purity [%] | RP-HPLC retention time | gradient | MW (kDa) | observed ions |
|---|---|---|---|---|---|---|---|---|---|
| Tsp1a-IR800$_{K4}$ | IR800 | 780 nm | 800 nm | 42 | 98 | 21.6 min | 0-95% solvent B | 4369.70 | $[M+3H]^{3+}$, $[M+4H]^{4+}$ |
| Tsp1a-DY684$_{K4}$ | DY684 | 660 nm | 680 nm | 18 | 95 | 24.2 min | 0-95% solvent B | 4365.67 | $[M+2H]^{2+}$, $[M+3H]^{3+}$ |
| Tsp1a-JA669$_{K4}$ | Janelia669 | 660 nm | 670 nm | 40 | 97 | 24.3 min | 0-95% solvent B | 3966.62 | $[M+2H]^{2+}$, $[M+3H]^{3+}$, $[M+4H]^{4+}$ |
| Tsp1a-BO665$_{K4}$ | Bodipy665 | 660 nm | 690 nm | 30 | 95 | 25.8 min | 0-95% solvent B | 3916.70 | $[M+2H]^{2+}$, $[M+3H]^{3+}$, $[M+4H]^{4+}$ |
| Tsp1a-CY7.5$_{K4}$ | CY7.5 | 760 nm | 790 nm | 39 | 96 | 28.0 min | 0-95% solvent B | 4019.86 | $[M+3H]^{3+}$, $[M+4H]^{4+}$, $[M+5H]^{5+}$ |
| Tsp1a-IR800$_{P}$ | IR800 | 780 nm | 800 nm | 60 | 98 | 21.7 min | 0-95% solvent B | 4682.87 | $[M+3H]^{3+}$, $[M+4H]^{4+}$, $[M+5H]^{5+}$ |
| Tsp1a-DY684$_{P}$ | DY684 | 660 nm | 680 nm | 29 | 96 | 26.8 min | 0-95% solvent B | 4678.84 | $[M+3H]^{3+}$, $[M+4H]^{4+}$, $[M+5H]^{5+}$ |
| Tsp1a-JA669$_{P}$ | Janelia669 | 660 nm | 670 nm | 55 | 98 | 24.3 min | 0-95% solvent B | 4279.80 | $[M+3H]^{3+}$, $[M+4H]^{4+}$, $[M+5H]^{5+}$ |
| Tsp1a-BO665$_{P}$ | Bodipy665 | 660 nm | 690 nm | 41 | 95 | 25.7 min | 0-95% solvent B | 4229.88 | $[M+3H]^{3+}$, $[M+4H]^{4+}$ |
| Tsp1a-CY7.5$_{P}$ | CY7.5 | 760 nm | 790 nm | 55 | 98 | 28.1 min | 0-95% solvent B | 4333.03 | $[M+3H]^{3+}$, $[M+4H]^{4+}$, $[M+5H]^{5+}$ |
| Tsp1a-K4 | —- | 280 nm | —- | 15 | 96 | 21.1 min | 0-95% solvent B | 3388.50 | $[M+2H]^{2+}$, $[M+3H]^{3+}$, $[M+4H]^{4+}$ |
| Tsp1a-Pra0 | —- | 280 nm | —- | 20 | 98 | 21.1 min | 0-95% solvent B | 3483.52 | $[M+2H]^{2+}$, $[M+3H]^{3+}$, $[M+4H]^{4+}$ |

**Table S1.** Chemical and photophysical features of fluorescently labeled Tsp1a peptides, including absorbance and emission maxima, synthetic yields, RP-HPLC retention times, purity, and ions observed using LC-MS.

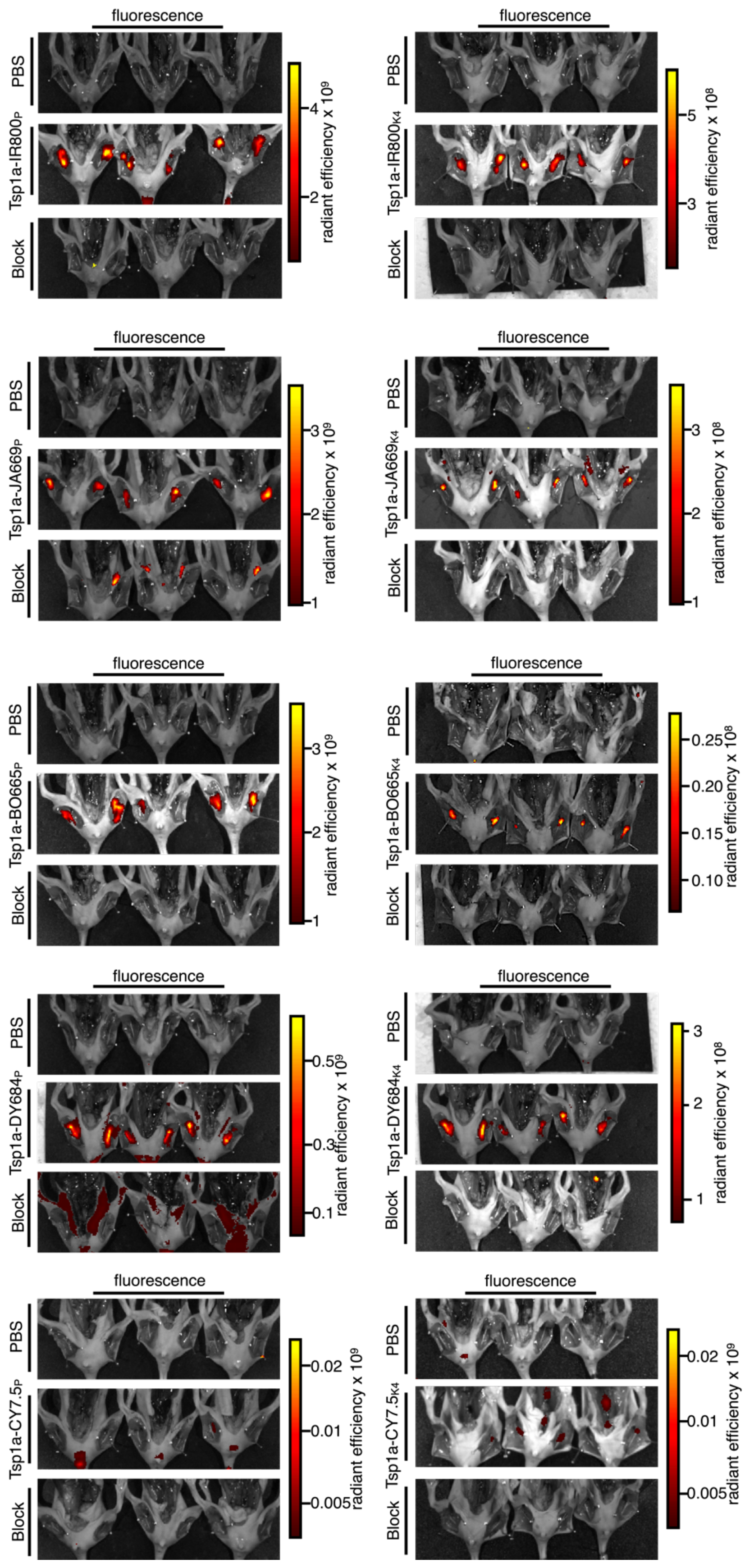
fluorescence
PBS
Tsp1a-IR800P
Block
radiant efficiency x 10^9
4
2
Tsp1a-IR800K4
radiant efficiency x 10^8
5
3
Tsp1a-JA669P
3
2
1
Tsp1a-JA669K4
Tsp1a-BO665P
Tsp1a-BO665K4
0.25
0.20
0.15
0.10
Tsp1a-DY684P
0.5
0.3
0.1
Tsp1a-DY684K4
Tsp1a-CY7.5P
0.02
0.01
0.005
Tsp1a-CY7.5K4

**Figure S9.** Pharmacokinetics of fluorescently labeled Tsp1a peptides. Epifluorescence images of animals injected with 100 μL PBS vehicle (top), Tsp1a tracer (1 nmol, 10 μM of tracer in 100 μL PBS), imaging agent (middle) or a Tsp1a-tracer/Tsp1a peptide block formulation (Tsp1a tracer, 10 μM, 1 nmol and Tsp1a peptide, 204 μM, 21 nmol in 100 μL PBS, bottom). Images of exposed peripheral nerves of mice were taken 30 min after tail vein injection.

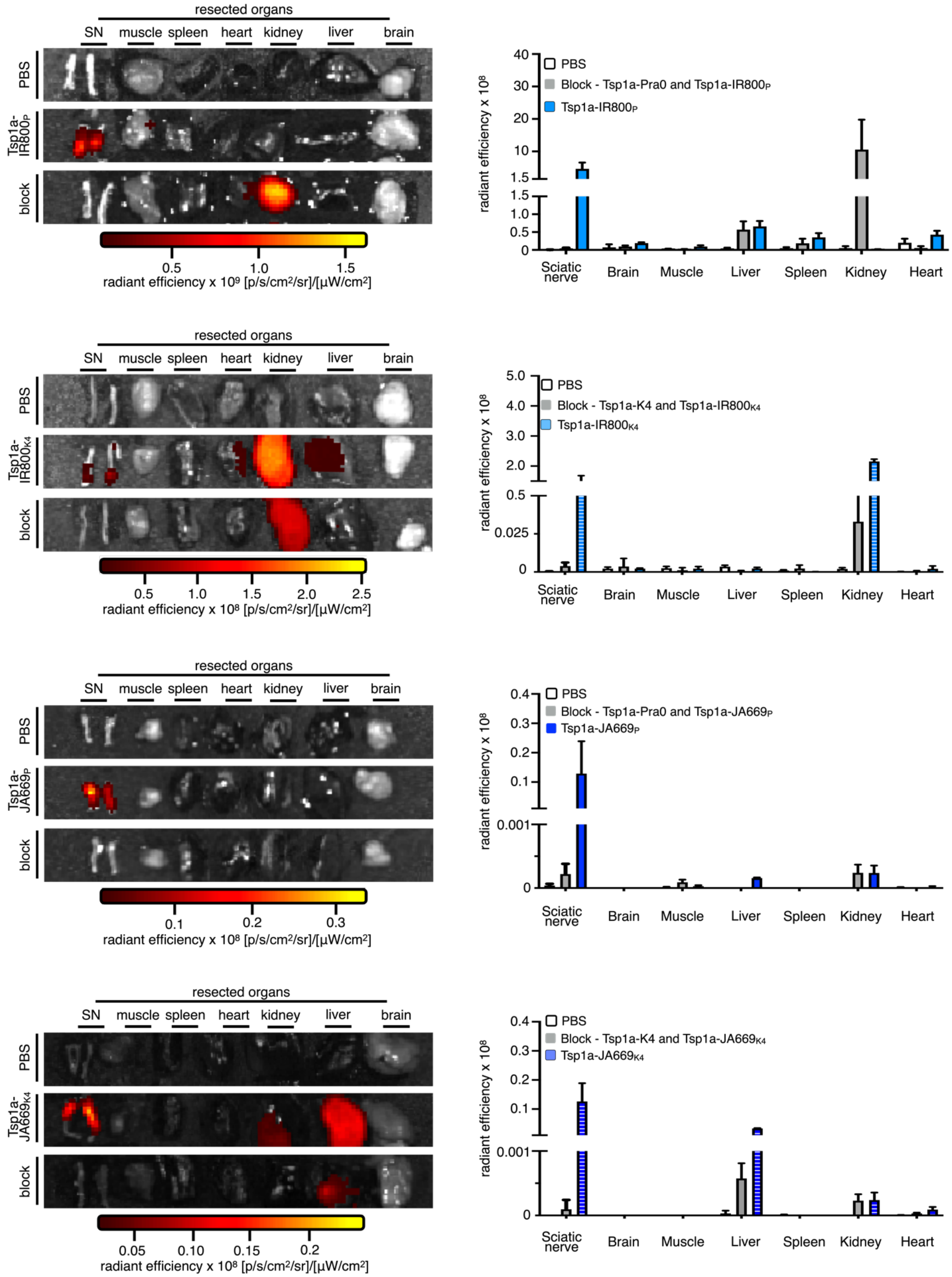

resected organs
SN
muscle
spleen
heart
kidney
liver
brain
PBS
Tsp1a-IR800P
block
0.5
1.0
1.5
radiant efficiency x 10⁹ [p/s/cm²/sr]/[µW/cm²]
radiant efficiency x 10⁸
PBS
Block - Tsp1a-Pra0 and Tsp1a-IR800P
Tsp1a-IR800P
40
30
20
10
1.5
1.5
1.0
0.5
0
Sciatic nerve
Brain
Muscle
Liver
Spleen
Kidney
Heart
resected organs
SN
muscle
spleen
heart
kidney
liver
brain
PBS
Tsp1a-IR800K4
block
0.5
1.0
1.5
2.0
2.5
radiant efficiency x 10⁸ [p/s/cm²/sr]/[µW/cm²]
radiant efficiency x 10⁸
PBS
Block - Tsp1a-K4 and Tsp1a-IR800K4
Tsp1a-IR800K4
5.0
4.0
3.0
2.0
0.5
0.025
0
Sciatic nerve
Brain
Muscle
Liver
Spleen
Kidney
Heart
resected organs
SN
muscle
spleen
heart
kidney
liver
brain
PBS
Tsp1a-JA669P
block
0.1
0.2
0.3
radiant efficiency x 10⁸ [p/s/cm²/sr]/[µW/cm²]
radiant efficiency x 10⁸
PBS
Block - Tsp1a-Pra0 and Tsp1a-JA669P
Tsp1a-JA669P
0.4
0.3
0.2
0.1
0.001
0
Sciatic nerve
Brain
Muscle
Liver
Spleen
Kidney
Heart
resected organs
SN
muscle
spleen
heart
kidney
liver
brain
PBS
Tsp1a-JA669K4
block
0.05
0.10
0.15
0.2
radiant efficiency x 10⁸ [p/s/cm²/sr]/[µW/cm²]
radiant efficiency x 10⁸
PBS
Block - Tsp1a-K4 and Tsp1a-JA669K4
Tsp1a-JA669K4
0.4
0.3
0.2
0.1
0.001
0
Sciatic nerve
Brain
Muscle
Liver
Spleen
Kidney
Heart

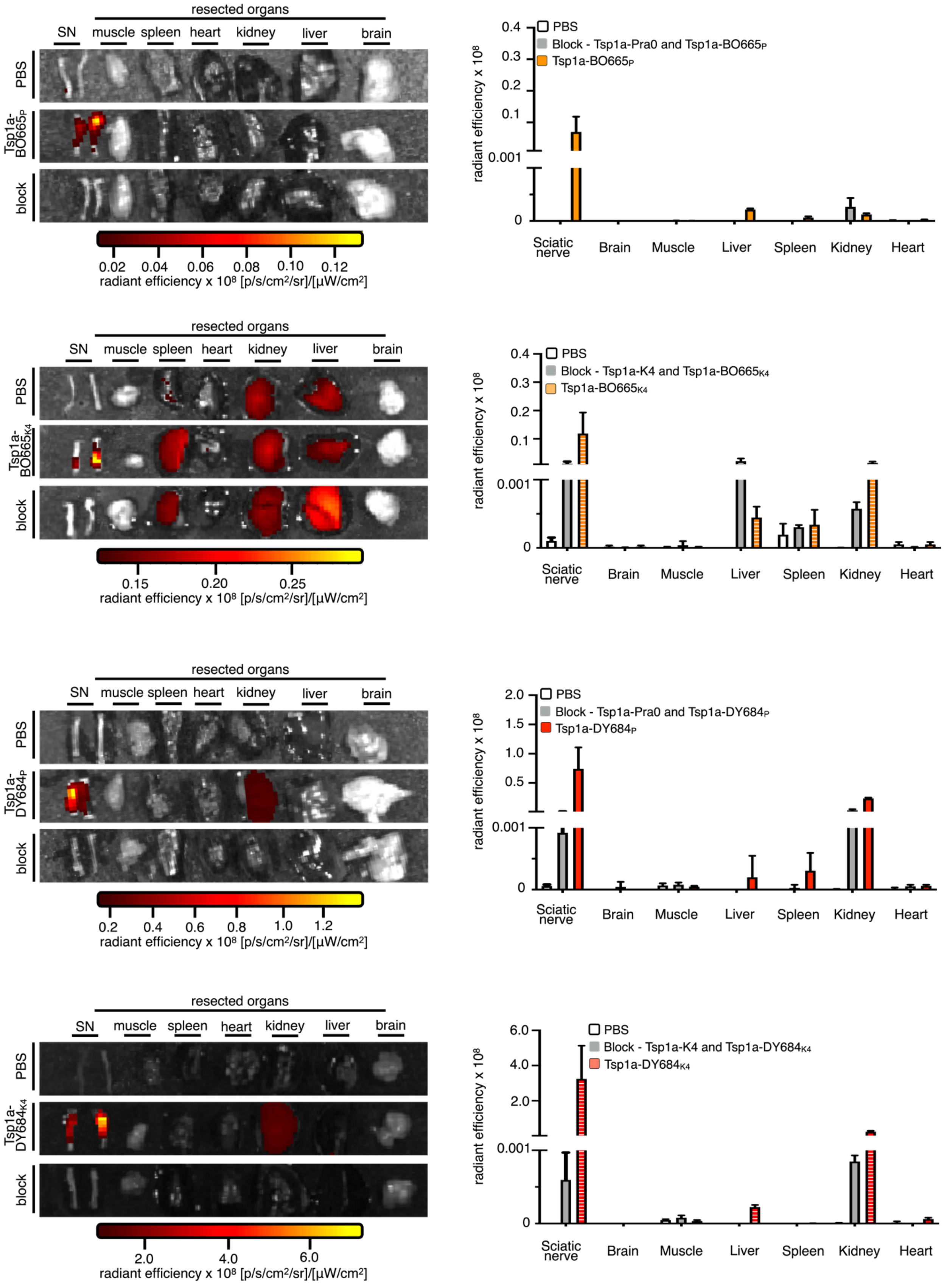
resected organs
SN
muscle
spleen
heart
kidney
liver
brain
PBS
Tsp1a-BO665P
block
0.02 0.04 0.06 0.08 0.10 0.12
radiant efficiency x 10⁸ [p/s/cm²/sr]/[µW/cm²]
radiant efficiency x 10⁸
PBS
Block - Tsp1a-Pra0 and Tsp1a-BO665P
Tsp1a-BO665P
0.4 0.3 0.2 0.1 0.001 0
Sciatic nerve
Brain
Muscle
Liver
Spleen
Kidney
Heart
resected organs
SN muscle spleen heart kidney liver brain
PBS
Tsp1a-BO665K4
block
0.15 0.20 0.25
radiant efficiency x 10⁸ [p/s/cm²/sr]/[µW/cm²]
radiant efficiency x 10⁸
PBS
Block - Tsp1a-K4 and Tsp1a-BO665K4
Tsp1a-BO665K4
0.4 0.3 0.2 0.1 0.001 0
Sciatic nerve Brain Muscle Liver Spleen Kidney Heart
resected organs
SN muscle spleen heart kidney liver brain
PBS
Tsp1a-DY684P
block
0.2 0.4 0.6 0.8 1.0 1.2
radiant efficiency x 10⁸ [p/s/cm²/sr]/[µW/cm²]
radiant efficiency x 10⁸
PBS
Block - Tsp1a-Pra0 and Tsp1a-DY684P
Tsp1a-DY684P
2.0 1.5 1.0 0.5 0.001 0
Sciatic nerve Brain Muscle Liver Spleen Kidney Heart
resected organs
SN muscle spleen heart kidney liver brain
PBS
Tsp1a-DY684K4
block
2.0 4.0 6.0
radiant efficiency x 10⁸ [p/s/cm²/sr]/[µW/cm²]
radiant efficiency x 10⁸
PBS
Block - Tsp1a-K4 and Tsp1a-DY684K4
Tsp1a-DY684K4
6.0 4.0 2.0 0.001 0
Sciatic nerve Brain Muscle Liver Spleen Kidney Heart

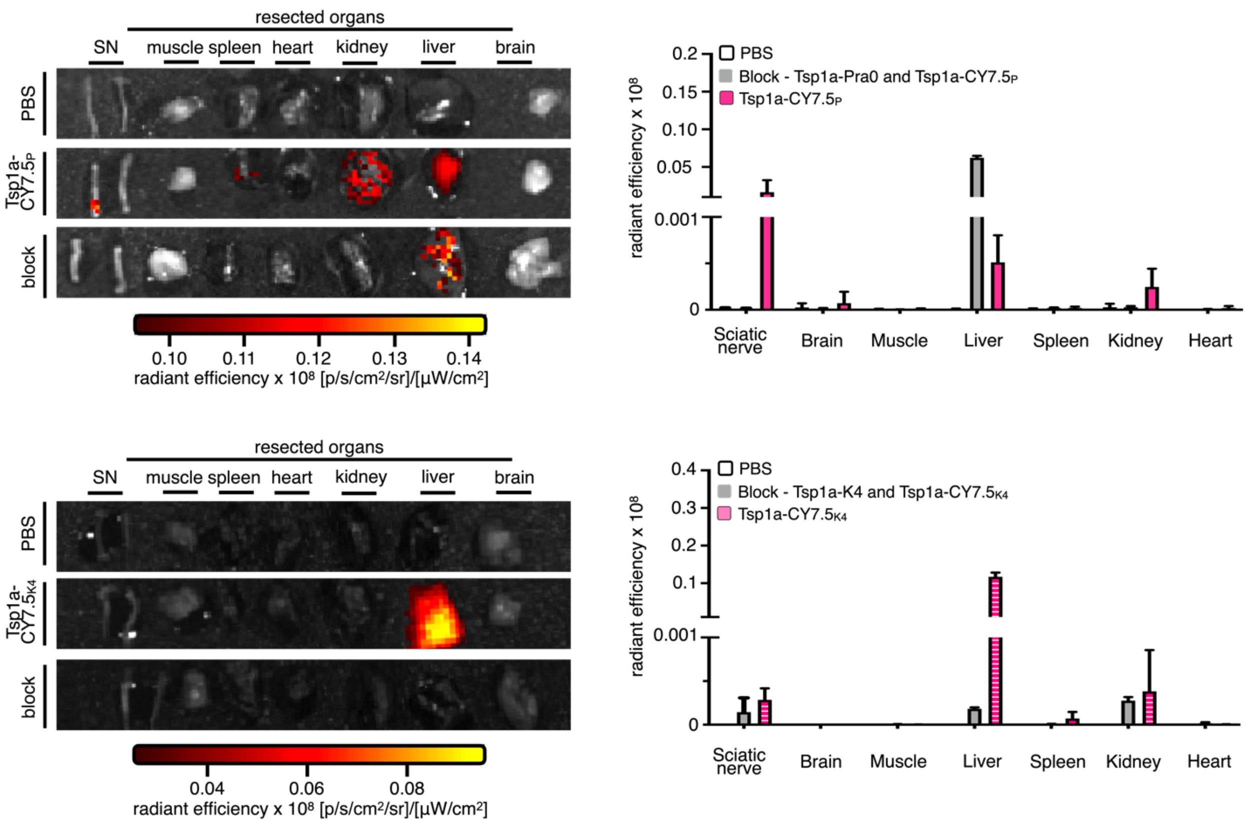


**Figure S10.** Pharmacokinetics of fluorescently labeled Tsp1a peptides. Epifluorescence images of resected right and left mouse peripheral nerves and corresponding organs that were injected with PBS, Tsp1a tracer or block formulation. High fluorescence intensities (due to fluorophore accumulation) were observed in sciatic nerves injected with Tsp1a tracers. No fluorescence was observed after 30 min in mice injected with vehicle or block formulation, except for Tsp1a-K4, which showed fluorescence in the kidneys, and signals in the liver for mice injected with Tsp1a-JA669$_{K4}$, Tsp1a-BO665$_{K4}$ and Tsp1a-CY7.5$_{K4}$. Right-hand panels show quantification of fluorescence in animals injected with PBS, Tsp1a-IR800$_{P}$ and block formulation. Statistics were calculated using a nonparametric Student's *t*-test. *P < 0.05; **P < 0.01; ***P < 0.001; ****P < 0.0001.

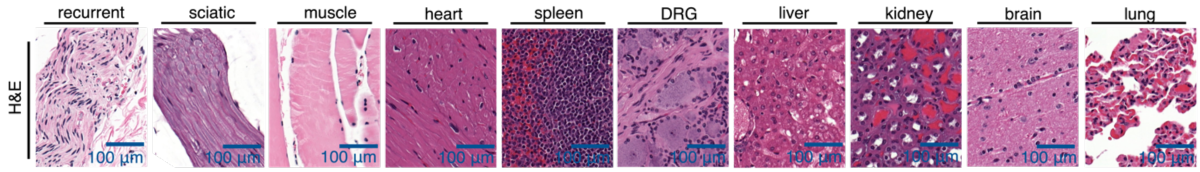


**Figure S11.** Slices from hematoxylin and eosin (H&E) staining of resected NHP peripheral nerves and organs. The topology of the peripheral nerves is similar to mouse and human samples.

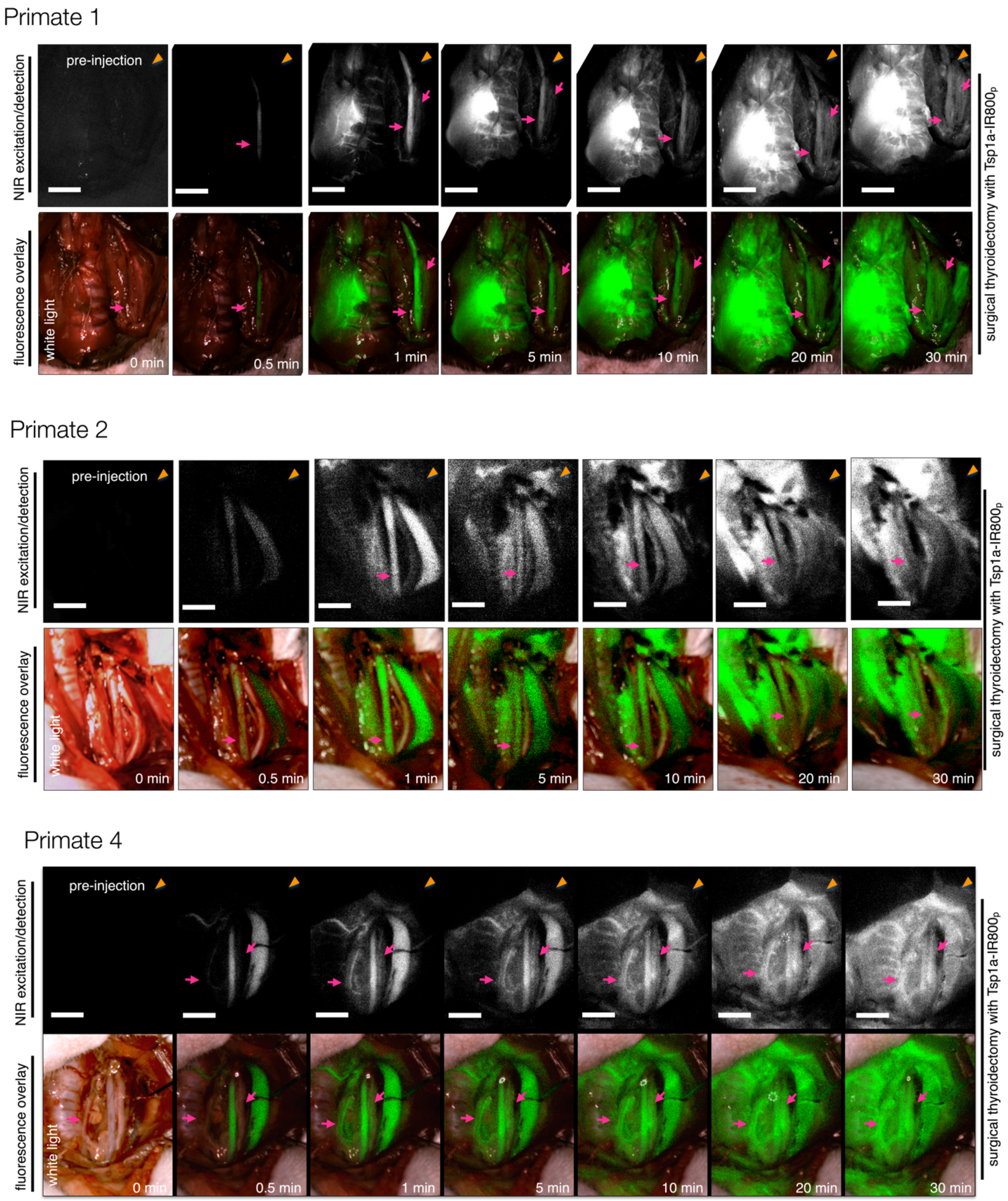
Primate 1
NIR excitation/detection
pre-injection
fluorescence overlay
white light
0 min
0.5 min
1 min
5 min
10 min
20 min
30 min
surgical thyroidectomy with Tsp1a-IR800p
Primate 2
NIR excitation/detection
pre-injection
fluorescence overlay
white light
0 min
0.5 min
1 min
5 min
10 min
20 min
30 min
surgical thyroidectomy with Tsp1a-IR800p
Primate 4
NIR excitation/detection
pre-injection
fluorescence overlay
white light
0 min
0.5 min
1 min
5 min
10 min
20 min
30 min
surgical thyroidectomy with Tsp1a-IR800p

**Figure S12.** In vivo performance of Tsp1a-IR800$_P$ in a clinical translatable setting. Fluorescence and overlay images from four NHPs subjected to a thyroidectomy after intravenous administration of Tsp1a-IR800$_P$ (250 μg kg$^{-1}$, in 5 mL PBS, delivered over ~60 s). Images of peripheral nerves highlight the recurrent laryngeal nerve (pink arrows) followed by demarcation of the vagus nerve over 30 min. Peripheral nerves are labeled (yellow arrows).

Primate 1

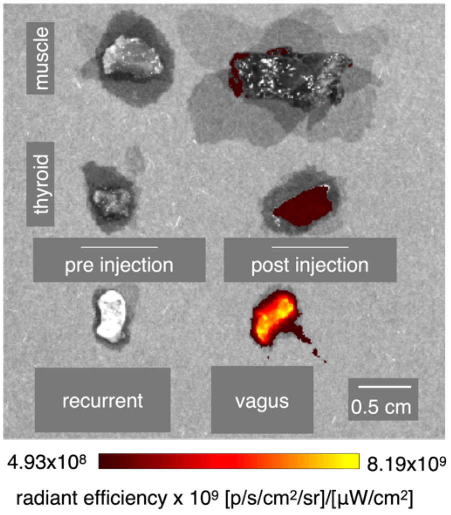


Primate 2

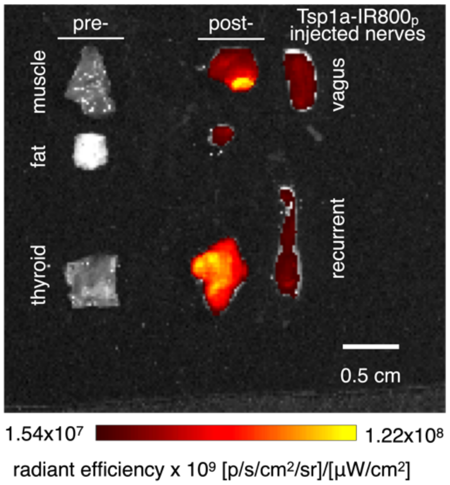


Primate 3

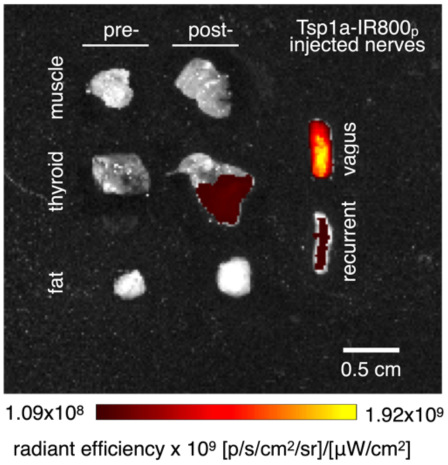

**Figure S13.** Ex vivo epifluorescence imaging of resected nerves and organs from primates that received intravenous Tsp1a-IR800$_P$ (250 μg kg$^{-1}$ in 5 mL PBS) prior to thyroidectomy. High fluorescence intensities (due to Tsp1a tracer accumulation) were observed in the peripheral nerves.
